# In vivo perturb-seq of cancer and microenvironment cells dissects oncologic drivers and radiotherapy responses in glioblastoma

**DOI:** 10.1101/2023.09.01.555831

**Authors:** S. John Liu, Joanna Pak, Christopher Zou, Timothy Casey-Clyde, Ashir A. Borah, David Wu, Kyounghee Seo, Thomas O’Loughlin, Daniel A. Lim, Tomoko Ozawa, Mitchel S. Berger, William A. Weiss, David R. Raleigh, Luke A. Gilbert

## Abstract

Genetic perturbation screens with single cell readouts have enabled rich phenotyping of gene function and regulatory networks. These approaches have been challenging *in vivo,* especially in adult disease models such as cancer, which include mixtures of malignant and microenvironment cells. Glioblastoma (GBM) is a fatal cancer, and methods of systematically interrogating gene function and therapeutic targets *in vivo*, especially in combination with standard of care treatment such as radiotherapy, are lacking. Here, we iteratively develop a multiplex *in vivo* perturb-seq CRISPRi platform for single cell genetic screens in cancer and tumor microenvironment cells that leverages intracranial convection enhanced delivery (CED) of sgRNA libraries into models of GBM. Our platform enables potent silencing of drivers of *in vivo* growth and tumor maintenance, as well as genes that sensitize GBM to radiotherapy. We find radiotherapy rewires transcriptional responses to genetic perturbations in an *in vivo* dependent manner, revealing heterogenous patterns of treatment sensitization or resistance in GBM. Furthermore, we demonstrate targeting of genes that function in the tumor microenvironment, enabling alterations of ligand-receptor interactions between immune/stromal cells following *in vivo* CRISPRi perturbations. In sum, we demonstrate the utility of multiplexed perturb-seq for *in vivo* single cell dissection of adult cancer and normal tissue biology across multiple cell types in the context of therapeutic intervention, a platform with potential for broad application.

## Background

Functional understanding of genes under physiological and disease states *in vivo* has been limited by a lack of approaches for multiplexed genetic perturbation at single cell resolution. CRISPR functional genomics have transformed our understanding of how genetic and epigenetic perturbations impact cell types and cell states^1^. However, existing functional genomic approaches *in vivo* have largely focused on population averaged phenotypes, obscuring cellular heterogeneity and cell-cell interactions that are critical for multicellular biology, disease, and response to stimuli^2–6^. Coupling CRISPR screening with single cell transcriptomics (perturb-seq) enables phenotyping of gene function using high dimensional gene expression readouts at the single cell level^7–13^. Perturb-seq phenotypes have been used to cluster genes by shared functions into pathways, allowing identification and characterization of genes with previously unknown function, and also to map epistatic relationships between genes. The single cell nature of perturb-seq also enables measurement of cellular heterogeneity, deconvolution of cell cycle effects, and is amenable to analysis of rare cell types. However, the majority of such efforts have focused on gene function *in vitro*^7–9,11^, utilize *ex vivo* perturbations^14,15^, or have been limited to developmental disorders^16,17^. As tissues and organs are comprised of numerous cell types that share physical and paracrine interactions, all while being influenced by the immune system, vasculature, and tissue architecture^18,19^, single cell transcriptomic screening using perturb-seq is especially warranted *in vivo*.

To establish an *in vivo* perturb-seq platform for multiplex interrogation of diseased or normal cells, we focused on glioblastoma (GBM), the most common primary malignant brain tumor^20^. GBM is incurable and exhibits remarkable cellular, genetic, and epigenetic heterogeneity^21,22^. GBM cells exist in multiple interchangeable cellular states^21,23^ embedded within an immunosuppressive microenvironment^24–26^, and GBM cultures *in vitro* do not adequately reflect the heterogeneity and dynamics of *in vivo* tumors^27^. Radiotherapy is the most effective adjuvant treatment for GBM, yet these tumors almost always recur^28–32^. Therefore, improved treatments for GBM are necessary, as well as strategies for enhancing the efficacy of standard of care therapies such as radiotherapy. Therefore, accurate functional genomics investigations of GBM as well as other solid malignancies compel the use of *in vivo* models.

Here, we iteratively developed a platform for *in vivo* perturb-seq in malignant and normal cells in the tumor microenvironment using convection enhanced delivery (CED), a technique that exploits bulk flow kinetics for enhanced delivery of viral vectors^33,34^. We demonstrate that transcriptomic phenotypes of oncogenic drivers can be defined in GBM malignant cells using perturb-seq, initially in cultures and then in orthotopic allografts. When combined with radiotherapy, CED perturb-seq reveals how radiotherapy treatment of established tumors rewires transcriptional responses to genetic perturbations. Furthermore, we demonstrate this platform allows for interrogation of cellular interactions between different cell types present in the tumor microenvironment upon genetic perturbation, highlighting the utility of perturbing cells in their intact environment.

## Results

### Syngeneic GBM tumors enable modeling of radiotherapy response and identification of oncogenic drivers

To begin to establish *in vivo* perturb-seq in tumor models with intact tumor immune microenvironments, we first utilized the GBM model GL261, which could model human tumors as well as be developed for CRISPRi loss of function screening to nominate candidates for further investigation using perturb-seq (Fig. 1A). The GL261 allograft is a well-established model that mimics aggressive human GBM with tumor features that include invasive growth, neovascularization, and orthotopic tumor formation with an intact immune system^35–37^. GL261 cells transplanted into the striata of C57BL/6 mice established tumors, and treatment with fractionated radiotherapy (RT, 2 Gy x 5 daily fractions) prolonged survival of GL261 allografts (Fig. 1B), consistent with the known efficacy but non-curative nature of radiotherapy in this model^38^, and more broadly in GBM. We then generated a GL261 cell line expressing CRISPRi machinery (dCas9-KRAB). To nominate genes that modulate GBM tumorigenesis and cellular responses to radiotherapy, we then performed large-scale CRISPRi screens in these cells *in vitro* against 5,234 cancer related and/or druggable genes^39^, in the presence or absence of radiotherapy (2 Gy x 5 daily fractions). We identified 230 genes modifying cell growth under control conditions (negative growth hits) and 49 genes modifying radiation resistance/sensitivity (Fig. 1C, S1A-B, Table S1). Gene ontology analysis of negative growth hits revealed enrichment for cell cycle, proteasome, and DNA replication genes, while radiation sensitizing hits revealed enrichment for non-homologous end joining, homologous recombination, and Fanconi anemia pathways (Fig. 1D, S1C). Internally controlled growth assays validated the phenotypes of 5 growth hits and 4 radiation sensitizing hits, demonstrating that our CRISPRi screening data effectively nominates genes of interest in the GL261 GBM model (Fig. S1D).

**Figure 1.**
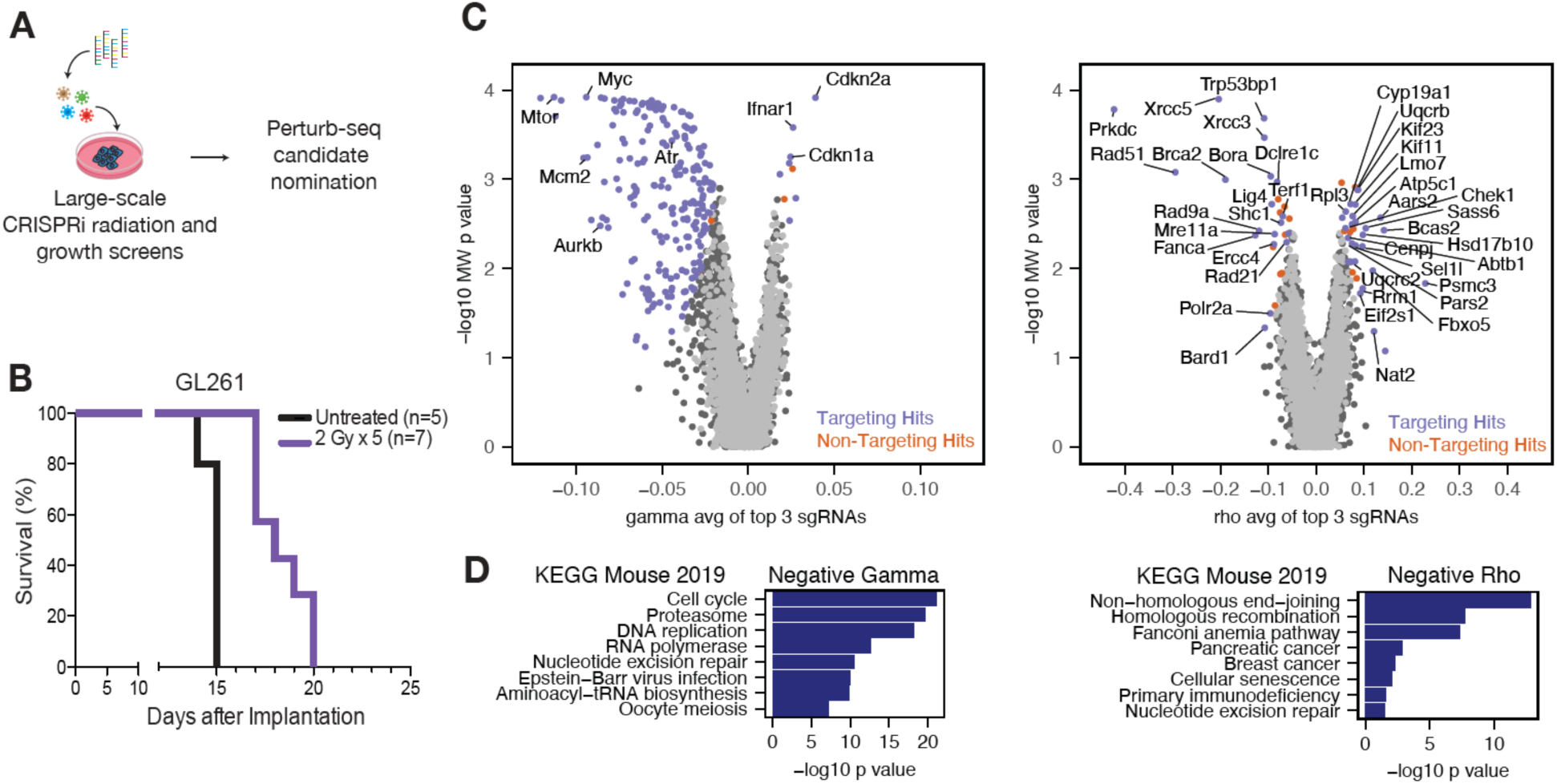
CRISPRi screens identify oncogenic drivers and modifiers of radiotherapy response in GBM. **A)** Schematic of *in vitro* screen workflow to nominate candidates for perturb-seq. **B)** Kaplan-Meier survival curves of GL261 intracranial tumor models treated with either fractionated radiotherapy (2 Gy x 5 fractions) to the whole brain or no treatment. **C)** Volcano plot of normalized growth (gamma) or radiation:growth ratio (rho) phenotypes in GL261 cells. A discriminant threshold score of 7 was used for both screen analyses to nominate hit genes for *in vivo* perturb-seq. **D)** Gene ontology enrichment analysis for negative gamma (left) and negative rho (right) screen hits identified in **(C)**.

### In vitro perturb-seq against oncogenic drivers and modifiers of radiotherapy response

To further characterize the functions of genes identified in the CRISPRi screens that regulate cell growth and radiotherapy response, we selected 48 hit genes to be further characterized by perturb-seq in GL261 cells grown *in vitro* (Fig. 2A). The top two sgRNAs (based on CRISPRi screen phenotypes) targeting each gene were cloned into dual sgRNA lentivirus vectors^40^, and a lentivirus pool containing this library and negative control sgRNAs was transduced at an MOI of ∼0.1 in GL261 cells stably expressing dCas9-KRAB. For *in vitro* perturb-seq, cultures were FACS sorted for sgRNA+ cells, treated with either fractionated radiotherapy (2 Gy x 5) or no treatment (0 Gy), and then harvested for scRNA-seq with direct capture of sgRNA tags^10^. As anticipated, sgRNA+ cells predominantly expressed two sgRNAs, with a small fraction of cells expressing four sgRNAs, reflective of cells infected by two lentiviral infection events (Fig. S2A). Nonetheless, the total sgRNA UMI counts in the vast majority of cells were equivalent to the UMI counts from the expected sgRNA A and sgRNA B targeting each gene (Fig. S2B). Only cells expressing the correct dual sgRNA vector were retained for further analysis, yielding a total of 8257 cells across two replicates for each treatment condition (Fig. S2C,D). As CRISPRi silences genes through transcriptional repression, we quantified the strength of gene silencing in cells pseudobulked by sgRNA identity and revealed 88.2% and 83.3% median target gene repression in the no treatment and radiotherapy conditions, respectively (Fig. 2B). Differential expression analysis revealed 17 target genes with greater than 100 differentially expression genes (adjusted p value < 0.05 and log2 fold change magnitude > 0.1) in the absence of radiotherapy (Fig. S2E). In the presence of radiotherapy, 8 gene targets passing quality control yielded greater than 100 differentially genes, when internally normalized to irradiated cells expressing non-targeting control sgRNAs (Fig. S2E). This pattern of differential gene expression changes was observed even after normalizing for differences in sgRNA coverage within each perturbation (Fig. S2F).

**Figure 2.**
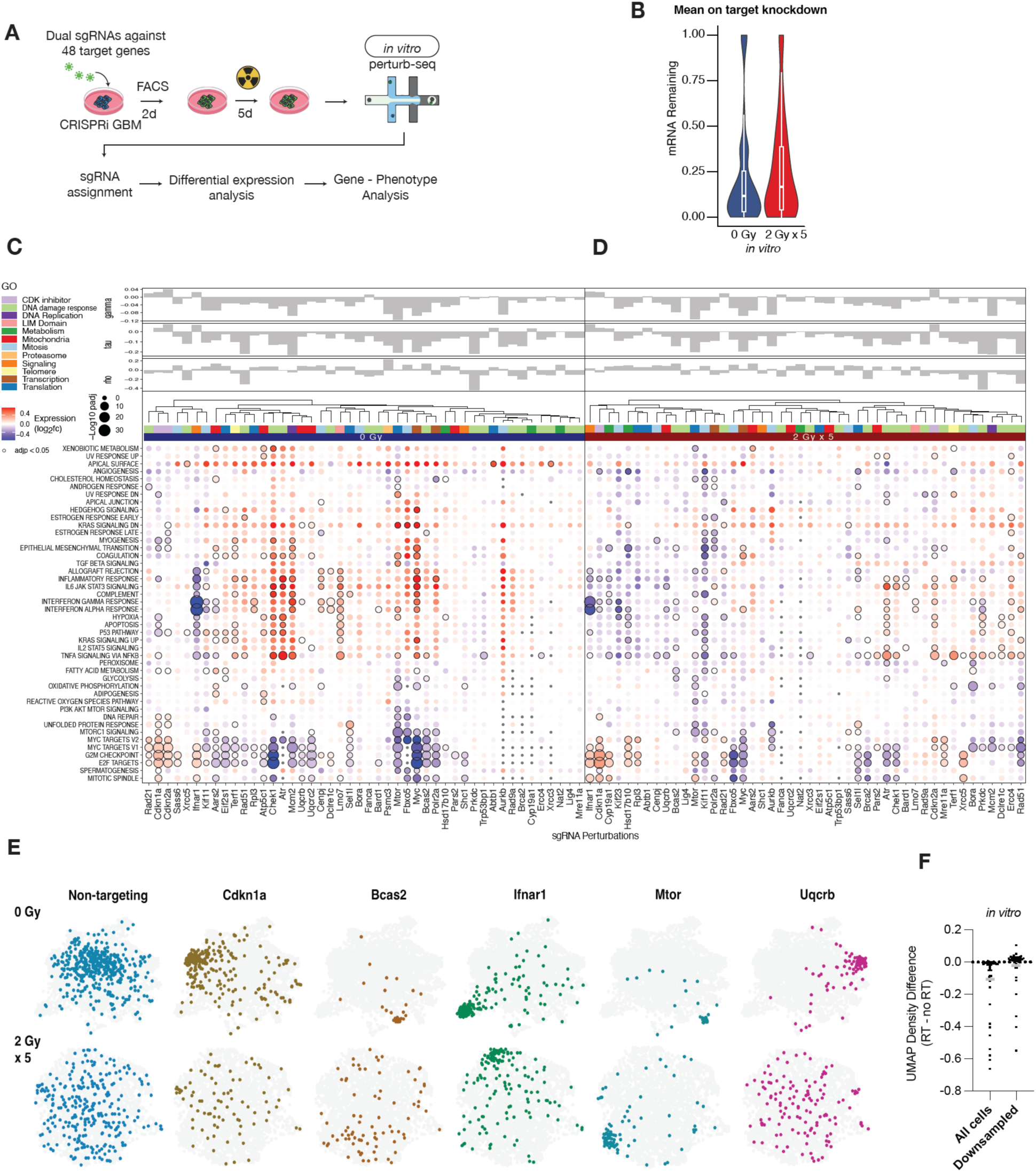
*In vitro* perturb-seq against oncogenic drivers and modifiers of radiotherapy response in GBM cells. **A)** Schematic of *in vitro* perturb-seq workflow against target genes nominated by genome-scale CRISPRi screens, combined with treatment with or without 2 Gy x 5 fractions of radiotherapy. **B)** Distribution of mean on-target knockdown levels for all *in vitro* perturbations in either treatment condition. Boxplots show 1st quartile, median, and 3rd quartile; whiskers represent 1.5 inter-quartile range. **C)** Bubble plot of gene set enrichment analyses of gene expression modules (rows) following sgRNA perturbations (columns) without radiotherapy. Expression represents log2 fold change normalized to non-targeting controls. Top bar charts show growth (gamma), radiation (tau), and radiation:growth ratio (rho) *in vitro* screen phenotypes. Gene ontology of perturbed target genes indicated. **D)** As in (C) but in radiotherapy conditions. Perturbation phenotypes were normalized to irradiated cells expressing non-targeting sgRNAs. **E)** LDA UMAP plots showing distribution of single cells expressing the indicated sgRNAs in either no radiotherapy (top) or radiotherapy (bottom) conditions. Cells which express sgRNAs other than the one highlighted in color are indicated in gray. **F)** Difference in UMAP gaussian kernel densities between perturbations in radiotherapy (RT) and no treatment (no RT) conditions. Gray bar = mean.

We then quantified transcriptomic phenotypes for each CRISPRi genetic perturbation using differential gene expression analysis followed by gene set enrichment analysis (Methods; Fig. 2C). Perturbation of 35 target genes in the absence of radiotherapy resulted in significant alteration (adjusted p value < 0.05) in at least one gene module (Fig. 2C). We observed expected changes for genetic perturbations known to be closely or causally annotated with specific gene modules. For example, the MTORC1 Signaling gene module was downregulated the most by perturbation of *Mtor*, while perturbation of *Myc* significantly downregulated MYC Targets V1 and V2 (Fig. 2C). Hierarchical clustering of perturbations revealed grouping by screen phenotypes, gene ontology of the target, and module alterations. For instance, positive growth hits *Cdkn1a, Cdkn2a*, and *Rad21* clustered together and were characterized by increased MYC Target, G2M Checkpoint, and E2F Target gene expression (Fig. 2C). Mitochondrial localized Complex III components *Uqcrb* and *Uqcrc2* clustered together, as did *Atr* and its downstream target *Chek1*. Despite different molecular functions, *Kif11* and *Ifnar1* clustered together based on significant downregulation of interferon α, interferon γ, and TNFα signaling, revealing convergent inflammatory responses from divergent population-based screen phenotypes (Fig. 2C).

To then define the molecular alterations induced by genetic perturbations in the context of radiotherapy, we quantified transcriptomic phenotypes of the same perturbations in GL261 cell cultures treated with fractionated radiotherapy (2 Gy x 5) (Fig. 2D). When internally normalized to irradiated cells expressing non-targeting control sgRNAs, regulators of DNA damage response (e.g. *Brca2, Ercc4, Lig4, Mre11a, Prkdc*), mitosis (e.g*. Bora*), and metabolism (e.g. *Hsd17b10, Cyp19a1*) exhibited more pathway alterations compared to the same perturbations in the absence of radiotherapy (Fig. 2D). In contrast to CRISPRi perturbations alone, certain perturbations in the context of radiotherapy such as *Myc, Atr, and Chek1* did not produce high magnitude expression changes in interferon/inflammatory responses or the p53 pathway (Fig. 2D). We therefore asked whether radiotherapy dominated the phenotypes of cells receiving combination genetic perturbation and radiotherapy by normalizing expression profiles with those of non-targeting sgRNA-expressing cells from unirradiated cultures. These gene expression changes demonstrated homogenous upregulation of Inflammatory Response (47 perturbations, adjusted p value < 0.05), apoptosis (30 perturbations, adjusted p value < 0.05), and downregulation of MYC Targets V2 (37 perturbations, adjusted p value < 0.05), and therefore radiotherapy *in vitro* potentially masks heterogeneity in molecular functions for genes that modify radiotherapy response (Fig. S2G). Consistent with this finding, linear discriminant analysis (LDA) revealed clustering of single cells by perturbations with distinct phenotypes in UMAP space, whereas single cells were more broadly distributed with the addition of radiotherapy (Fig. 2E). UMAP densities for perturbed cells were lower in radiotherapy conditions, even after normalizing for cell coverage by downsampling to equal numbers of cells in radiotherapy and no radiotherapy conditions for each perturbations (Fig. 2F).

### In vivo perturb-seq against oncogenic drivers and modifiers of radiotherapy response

Glioblastoma cells are generally more resistant to radiotherapy *in vivo* than *in vitro*^41^, likely due to multiple tumor-specific factors that cannot be recapitulated *in vitro*^21,42^. To define the molecular pathways underlying oncogenic drivers and radiotherapy response in GBM, we established an *in vivo* perturb-seq platform in GL261 cells by transducing cells *ex vivo* with sgRNA libraries prior to *in vivo* tumorigenesis (hereafter referred to as *in vivo* pre-infected perturb-seq) (Fig. 3A). A GFP-tagged dual sgRNA lentivirus library targeting the same 48 genes from perturb-seq *in vitro* was transduced into GL261 dCas9-KRAB cell cultures, which were puromycin selected for sgRNA expression, and then transplanted intracranially into C57BL/6 mice. Tumors were allowed to expand for 5 days and were then treated with fractionated radiotherapy (2 Gy x 5) or no treatment (0 Gy), followed by single cell dissociation and scRNA-seq with direct capture of sgRNA tags^10^. GFP sorted cells as well as unsorted tissue dissociates were processed to capture the full extent of cellular heterogeneity. 20 distinct stromal or immune microenvironment cell types were identified based on transcriptomic profiles, in addition to GL261 malignant cells, which were readily identified by mRNA expression of BFP (CRISPRi marker) and sgRNAs (Fig. 3B,C, S3A). As was observed for *in vitro* perturb-seq, sgRNA+ cells predominantly expressed two sgRNAs (Fig. S3B), and the total sgRNA UMI’s were equivalent to the UMI counts from the expected sgRNA A and sgRNA B targeting each gene (Fig. S3C). 30,878 *in vivo* GL261 cells expressing both expected sgRNAs for each gene target were identified across biological triplicate experiments from no radiotherapy or radiotherapy conditions (Fig. S3D, E). Analysis of CRISPRi knockdown revealed 96.5% and 92.2% median target gene repression in the no treatment and radiotherapy conditions, respectively (Fig. 3D).

**Figure 3.**
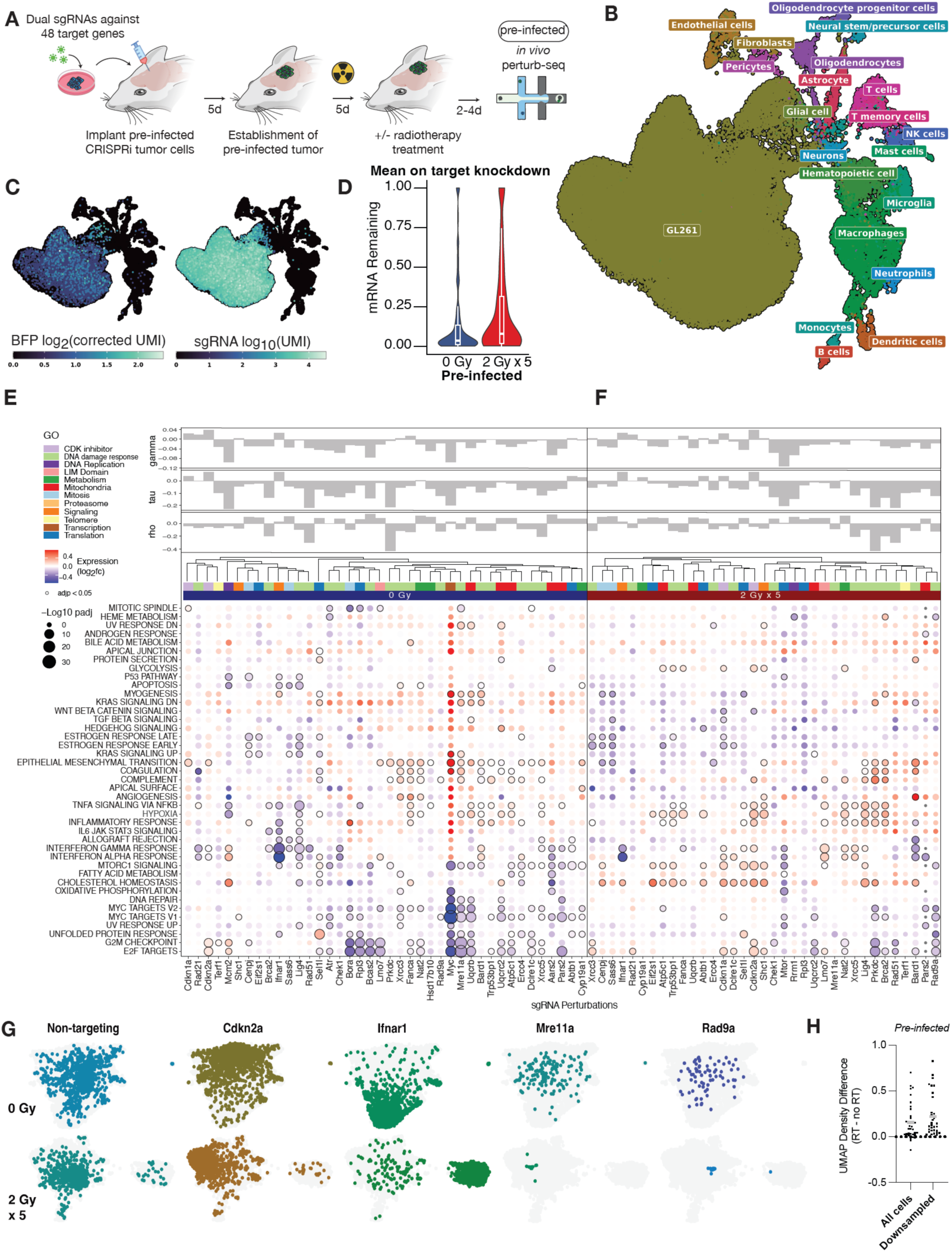
*In vivo* perturb-seq against oncogenic drivers and modifiers of radiotherapy **response using pre-infected GL261 cells. A)** Schematic of *in vivo* pre-infected perturb-seq workflow against target genes nominated by large scale CRISPRi screens, combined with treatment with or without 2 Gy x 5 fractions of radiotherapy. **B)** Integrated scRNA-seq UMAP of malignant and stromal/microenvironment cells from orthotopic tumors containing pre-infected GL261 cells, including sorted and unsorted cells. **C)** Integrated scRNA-seq UMAP from (B) overlayed with BFP (marker for dCas9-KRAB; left) or sgRNA (right) expression levels. **D)** Distribution of mean on-target knockdown levels for all *in vivo* pre-infected perturbations in either treatment condition. Boxplots show 1st quartile, median, and 3rd quartile; whiskers represent 1.5 inter-quartile range. **E)** Bubble plot of gene set enrichment analyses of gene expression modules (rows) following sgRNA perturbations (columns) without radiotherapy for *in vivo* pre-infected perturb-seq. Expression represents log2 fold change normalized to non-targeting controls. Top bar charts show growth (gamma), radiation (tau), and radiation:growth ratio (rho) *in vitro* screen phenotypes. Gene ontology of perturbed target genes indicated. **F)** As in (E) but in radiotherapy conditions. Perturbation phenotypes were normalized to cells from irradiated tumors that expressed non-targeting sgRNAs. **G)** LDA UMAP plots showing distribution of single cells from *in vivo* pre-infected experiments expressing the indicated sgRNAs in either no radiotherapy (top) or radiotherapy (bottom) conditions. Cells which express sgRNAs other than the one highlighted in color are indicated in gray. H) Difference in UMAP gaussian kernel densities between perturbations in radiotherapy (RT) and no treatment (no RT) conditions. Gray bar = mean.

We then generated perturbation-phenotype maps using differential gene expression analysis (Methods), revealing 24 target genes with greater than 100 differentially expression genes (adjusted p value < 0.05 and log2 fold change magnitude > 0.1) in the absence of radiotherapy and 34 with radiotherapy (normalized to non-targeting sgRNA in radiotherapy conditions) (Fig. S3F). The increased number of differentially expressed genes in the radiotherapy conditions was observed after normalizing for sgRNA coverage per perturbation (Fig. S3G). Perturbation of 40 and 30 target genes passing quality control filters resulted in significant alteration (adjusted p value < 0.05) in at least one gene module in the absence or presence of radiotherapy, respectively (Fig. 3E,F). Our analysis of gene module alterations in the absence of radiation demonstrated expected biology. For example, the MYC Targets V2 gene module was downregulated the most by perturbation of *Myc*, while interferon and inflammatory pathways were downregulated the most by perturbation of *Ifnar1* (Fig. 3E). Surprisingly, perturbation of certain genes with radiation sensitizing CRISPRi screen phenotypes, such as *Mre11a, Fanca*, and *Ercc4*, exhibited significant gene modules alterations even in the absence of radiotherapy *in vivo* (Fig. 3E), an observation that was not seen from *in vitro* perturb-seq (Fig. 2C) despite adequate coverage of these perturbations in both experimental contexts (Fig. S2D, S3E). These data underscore the importance of *in vivo* functional interrogation of genes discovered from more traditional *in vitro* approaches.

We then analyzed gene module alterations induced by genetic perturbations in GL261 *in vivo* tumors that were treated with radiotherapy. A cluster of perturbations targeting genes involved in DNA damage response (*Xrcc5, Lig4, Prkdc, Brca2, Mre11a, Rad51*) exhibited radiotherapy-dependent upregulation of hypoxia and inflammatory response (Fig. 3F), while other genes with radiotherapy sensitizing phenotypes such as *Bard1* showed decreases in interferon α and interferon γ response. Cholesterol metabolism was significantly upregulated (adjusted p value < 0.05) in 15 perturbations in the context of radiotherapy, compared to 3 in the absence of radiotherapy (Fig. 3E, F). These pathway alterations suggest that radiotherapy rewires the transcriptional responses to loss of genes that regulate oncogenic function or radiation responses in GBM (Fig. 3F). These radiotherapy-dependent phenotypes were also evident in LDA UMAP space (Fig. 3G). Despite expected depletion of cells expressing sgRNAs against DNA repair factors such as *Mre11a* or *Rad9a* in radiotherapy conditions, the increase in UMAP density observed for cells perturbed in the context of radiotherapy was maintained even after controlling for cell counts (Fig. 3H). While *in vivo* pre-infected perturb-seq revealed many *in vivo-* and radiotherapy-dependent phenotypes, one limitation of this approach is the relatively low coverage of perturbations with strong negative growth phenotypes, which precluded analysis of critical gene targets such as *Mtor* or *Aars2* in the radiotherapy conditions (Fig. S3E).

### Convection enhanced delivery enables in vivo perturb-seq after tumor establishment

To enable single cell CRISPR screening in tumor cells genetically perturbed within their native physiologic context, we established *in vivo* perturb-seq using convection enhanced delivery (CED) to deliver sgRNA lentiviruses directly into orthotopic GBMs (Fig. 4A). We transplanted GL261 GBM cells stably expressing CRISPRi machinery (dCas9-KRAB) into the striata of C57BL/6 mice (Fig. S4A). Following tumor establishment, lentiviral dual-sgRNA libraries targeting the 48 genes nominated from *in vitro* CRISPRi screens (Fig. 1) and also targeted in *in vitro/pre-infected* perturb-seq (Fig. 2A, 3A), plus non-targeting controls were delivered to intracranial tumors using CED. Animals were treated with either cranial fractionated radiotherapy (2 Gy x 5 daily fractions) or no treatment (0 Gy). Tumors were then microdissected, dissociated, sorted for sgRNA+ cells, and harvested for scRNA-seq with direct capture of sgRNA tags^10^. Integration of FACS sorted sgRNA positive cells and sgRNA negative cells from unsorted populations revealed 22 immune, stromal, and malignant cell populations across UMAP space (Fig. 4B; S4B). Analysis of sgRNA UMI counts confirmed single lentivirus vector integration events, as total sgRNA UMI counts were highly correlated with the sum of sgRNA UMI counts from sgRNA A and sgRNA B (Pearson R = 1.00) (Fig. S4C-E). Nearly all sgRNA positive cells were GL261 malignant cells, and 2153 *in vivo* GL261 cells expressing only the expected dual sgRNA vectors were retained (5 pooled animals per condition-replicate) (Fig. 4C, D, Fig. S4F, G). In contrast to *in vivo* perturb-seq experiments using pre-infected cells, CED perturb-seq allowed better recovery of sgRNAs targeting essential genes such as *Myc* and *Mtor*, thus enabling transcriptional phenotyping of genes important in tumor maintenance (Fig. S4F). Analysis of CRISPRi knockdown efficacy revealed 95.5% and 95.6% median target gene repression in the no treatment and radiotherapy conditions, respectively (Fig. 4E).

**Figure 4.**
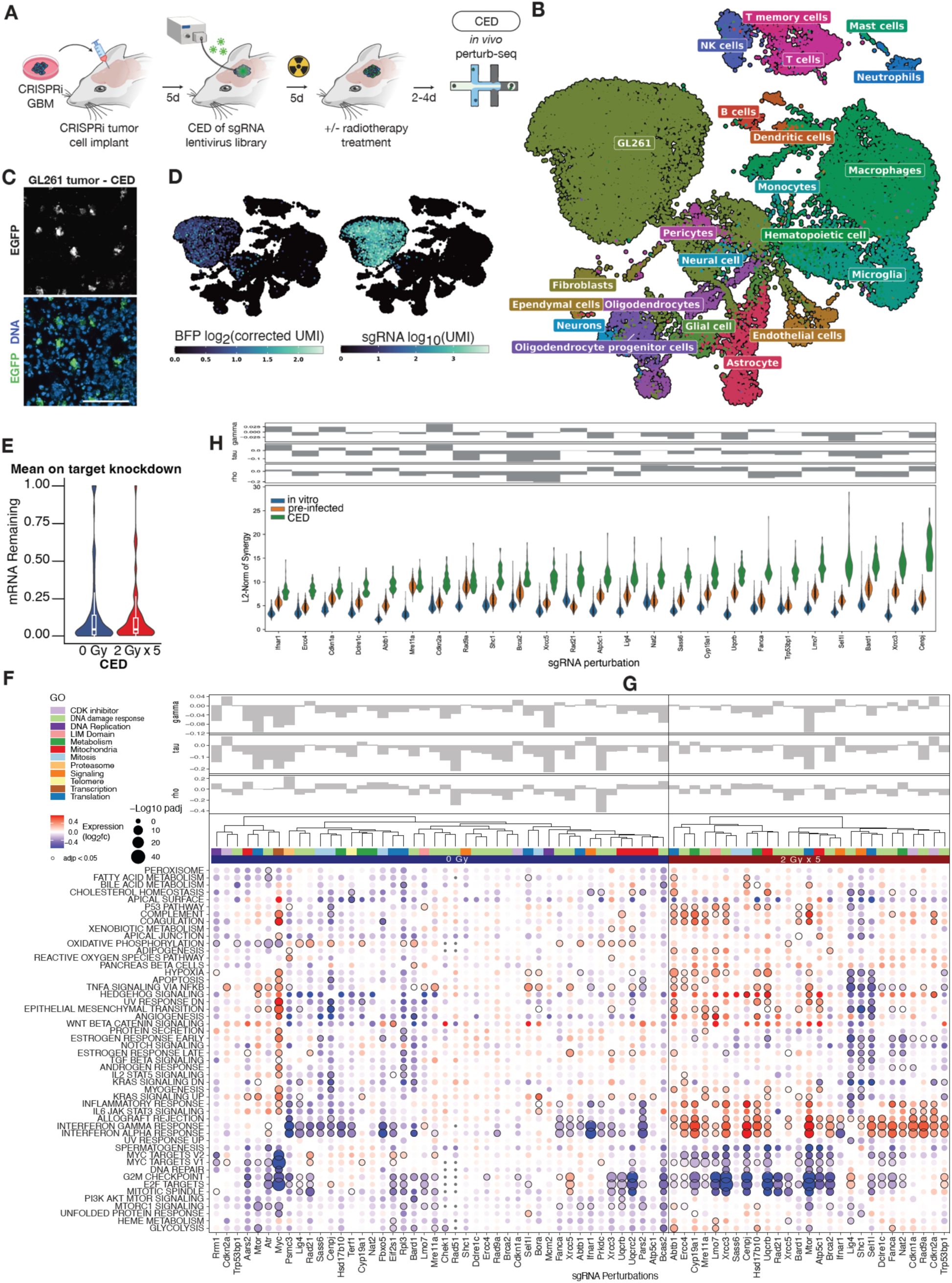
*In vivo* perturb-seq after tumor establishment using convection enhanced delivery. **A)** Schematic of *in vivo* CED perturb-seq workflow against target genes nominated by genome-scale CRISPRi screens, combined with treatment with or without 2 Gy x 5 fractions of radiotherapy. **B)** Integrated scRNA-seq UMAP of malignant and stromal/microenvironment cells from orthotopic GL261 tumors following CED, including sorted and unsorted cells. **C)** Confocal image of GL261 orthotopic tumor transduced with a sgRNA library tagged with EGFP. Scale bar, 100 µm. **D)** Integrated scRNA-seq UMAP of malignant and stromal/microenvironment cells from (B) overlayed with BFP (marker for dCas9-KRAB; left) or sgRNA (right) expression levels. **E)** Distribution of mean on-target knockdown levels for all *in vivo* CED perturbations in either treatment condition. Boxplots show 1st quartile, median, and 3rd quartile; whiskers represent 1.5 inter-quartile range. **F)** Bubble plot of gene set enrichment analyses of gene expression modules (rows) following sgRNA perturbations (columns) without radiotherapy for *in vivo* CED perturb-seq. Expression represents log2 fold change normalized to non-targeting controls. Top bar charts show growth (gamma), radiation (tau), and radiation:growth ratio (rho) *in vitro* screen phenotypes. Gene ontology of perturbed target genes indicated. **G)** As in (E) but in radiotherapy conditions. Perturbation phenotypes were normalized to cells from irradiated tumors that expressed non-targeting sgRNAs following CED. **H)** L2-norm synergy scores for each sgRNA perturbation across all experimental contexts (*in vitro*, *in vivo* pre-infected, *in vivo* CED).

We then quantified transcriptomic phenotypes for each CRISPRi genetic perturbation following CED using differential expression analysis (Methods), revealing 8 targets that passed quality and coverage filters with greater than 100 differentially expression genes (adjusted p value < 0.05 and log2 fold change magnitude > 0.1) in the absence of radiotherapy and 15 with radiotherapy (Fig. S4H). The increased number of differentially expressed genes in the radiotherapy conditions was observed after normalizing for sgRNA coverage for each perturbation (Fig. S4I). Perturbation of 38 and 26 target genes passing quality control filters resulted in significant alteration (adjusted p value < 0.05) in at least one gene module in the absence or presence of radiotherapy, respectively (Fig. 4F, G). In the absence of radiotherapy, as was observed for *in vitro* perturb-seq (Fig. 2C) and pre-infected perturb-seq (Fig. 3E), MTORC1 Signaling was downregulated the most by perturbation of *Mtor* via CED, and perturbation of *Myc* significantly downregulated MYC Targets V1 and V2 (Fig. 4F). Hierarchical clustering of gene expression changes caused by perturbations demonstrated multiple clusters which are distinguished by which gene modules were altered, or by the known functions of the target genes (Fig. 4F). While *in vitro* growth screen phenotypes in the absence of radiotherapy correlated with the clustering of many perturbations, *in vivo* transcriptional responses were in some cases shared despite opposing screen phenotypes *in vitro* (Fig. 4F). For example, *Ifnar1* perturbation downregulated interferon and inflammatory signaling, as did perturbation of *Lig4* or *Rad21*, which play roles in DNA damage response and show opposing phenotypes in growth screens compared to *Ifnar1* (Fig. 4F, Table S1). In contrast, perturbation of *Atr* resulted in upregulation of interferon responses, while perturbation of *Myc* upregulated STAT3/5 signaling in CED experiments (Fig. 4F). *Atr, Myc,* and *Mtor* perturbations were also characterized by shared downregulation of oxidative phosphorylation genes (Fig. 4F). Mitochondrial components (*Uqcrb, Uqcrc2, Pars2, Atp5c1*) clustered together as well.

We then analyzed CED perturbations in the context of *in vivo* cranial radiotherapy. Normalized to non-targeting sgRNAs in radiotherapy conditions, heterogenous alterations in gene expression modules were observed in radiotherapy conditions (Fig. 4G). In contrast to perturbations without radiotherapy, CED perturbations in the presence of radiotherapy exhibited upregulation of inflammatory/interferon signaling, as well as activation of p53 pathway, even after normalizing to cells from the same irradiated tumors that express non-targeting sgRNAs (Fig. 4G). Multiple clusters of perturbations were observed following hierarchical clustering, and these gene expression profiles spanned classes of transcriptomic alterations characterized by (1) low proliferative, low inflammatory, and low epithelial mesenchymal transition genes (*Sel1l, Lig4, Shc1*), (2) low proliferative and high inflammatory signaling (e.g. *Cyp19a1, Xrcc3, Cenpj, Mtor, Atp5c1, Brca2, Fanca*), and (3) variable proliferative and high inflammatory signaling (*e.g. Rad9a, Trp53bp1, Cdkn1a, Cdkn2a*). Perturbations from each of these classes demonstrated radiotherapy-dependent phenotypes, as evidenced by preferential aggregation of cells with perturbations such as *Ercc4, Fanca,* and *Shc1* in LDA UMAP space with radiotherapy (Fig. S5A, B). As was the case for pre-infected perturb-seq (Fig. 3H) (but not *in vitro* perturb-seq (Fig. 2F)), UMAP density was greater for CED perturbations combined with radiotherapy (Fig. S5C).

Given that *in vivo* perturbations rewired transcriptional responses following radiotherapy, we asked whether perturbations combined with radiotherapy could be modeled as a linear combination of perturbation effects in the absence of radiotherapy and radiotherapy effects alone, with potential for synergy. We used CINEMA-OT (causal independent effect module attribution with optimal transport)^43^ to quantify synergy scores, defined as the difference between the observed phenotype for combination treatment and the predicted phenotype assuming a linear additive relationship between genetic perturbation and radiotherapy. Synergy scores were greater for the majority of perturbations with CED compared to either *in vitro* or pre-infected contexts (Fig. 4H). *In vitro* CRISPRi screen phenotypes for radiation overlayed with the synergy metric showed little correlation (Fig. 4H), demonstrating the utility of *in vivo* CED perturb-seq to reveal the roles of radiation sensitizing and resistance gene targets *in vivo*.

### Convection enhanced delivery enables in vivo perturb-seq within the tumor microenvironmemnt

As cancers comprise both malignant and non-malignant host cells of the stroma or immune microenvironment, we asked whether *in vivo* CED perturb-seq could be applied toward stromal or immune microenvironment cells in GBM tumors. To that end, we used the SB28 syngeneic mouse GBM model, which is driven by *Nras* overexpression and establishes a myeloid cell-rich tumor immune microenvironment similar to human GBM^44,45^. Although both GL261 and SB28 form orthotopic tumors in immunocompetent animals, SB28 tumors harbor a dearth of T cells and resistance to immunotherapy that are more reflective of human GBM^46^. SB28 cells that did not express CRISPRi machinery were transplanted intracranially into immunocompetent mice constitutively expressing dCas9-KRAB^47^, enabling cell type-selective sgRNA stabilization and knockdown of genes in microenvironment cells but not in malignant cells (Fig. 5A). Target genes (*Apoe, C1qa, Cd44, Cd74, Lyz2, Ptprc*) were selected for perturbation based on cell type-specific expression or function in the myeloid lineage, given their importance in GBM pathogenesis and potential for therapeutic intervention^48,49^. We delivered a lentiviral sgRNA library (3 individual sgRNAs targeting each of 6 target gene, including a non-targeting sgRNA) by CED into SB28 tumors, dissected tumors 5 days following CED, and performed scRNA-seq with direct sgRNA capture of both FACS sorted and unsorted tumors. Analysis revealed a diversity of non-cancer cell types (e.g. macrophages, astrocytes, endothelial cells) within the tumor microenvironment, existing alongside SB28 cancer cells, which were readily distinguishable by *Nras* overexpression (Fig. 5B, D, Fig. S6,S7A). Consistent with their high abundance in SB28 allografts, macrophages, microglia, as well as glial cells represented the large majority of sgRNA-expressing cells in these tumors, spanning 6909 FACS sorted cells from quadruplicate experiments (Fig. 5C,D). Analysis of CRISPRi knockdown levels revealed heterogeneous target gene suppression across the different cell types analyzed in this experiment, and only sgRNAs with greater than 30% knockdown (less than 70% mRNA remaining) were retained for subsequent analysis in each cell type (Fig. S7B).

**Figure 5.**
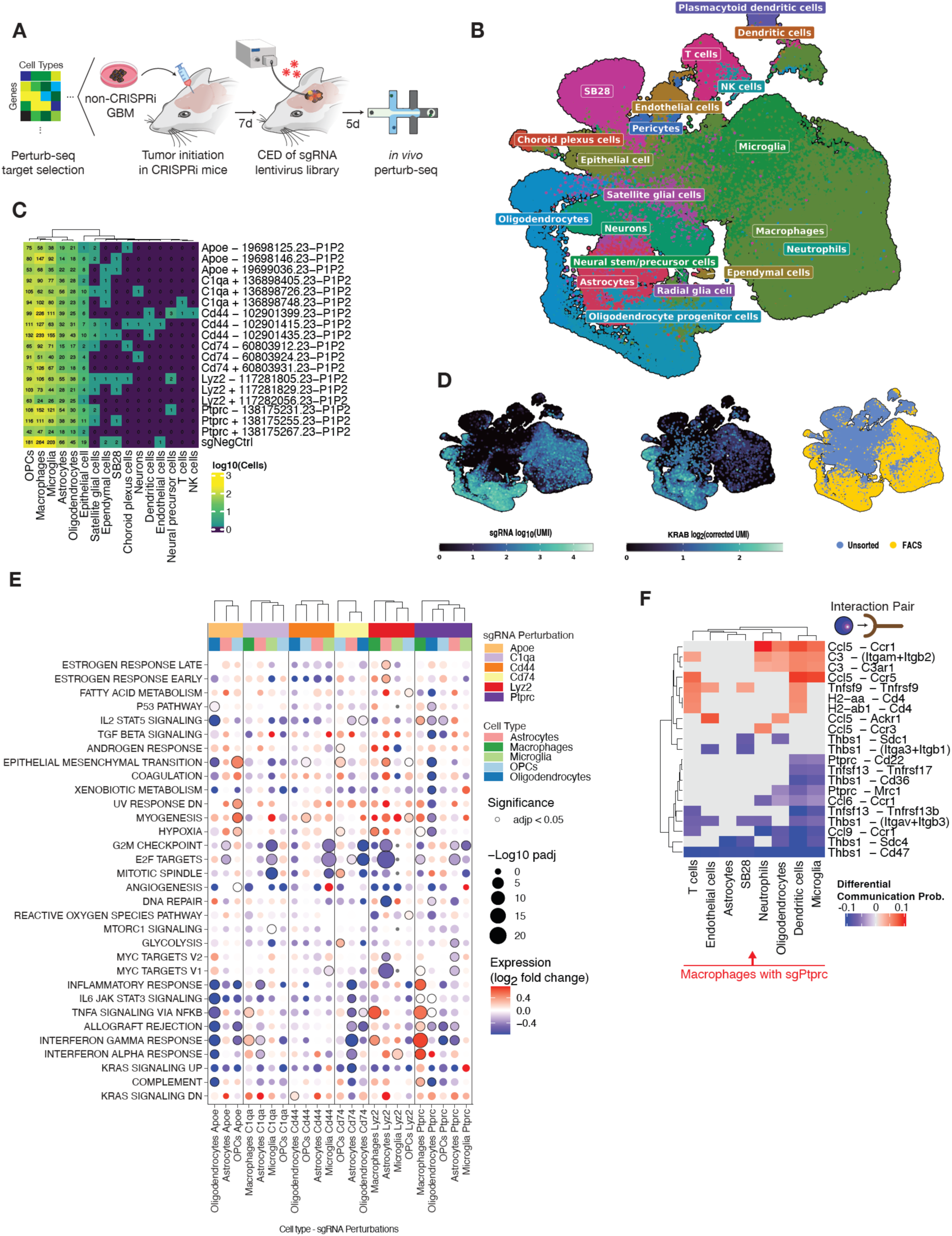
In vivo perturb-seq in tumor microenvironment cells. **A)** Schematic of *in vivo* CED perturb-seq in orthotopic SB28 GBM allograft models against target genes. **B)** Integrated scRNA-seq UMAP of malignant and stromal/microenvironment cells from orthotopic SB28 tumors following sgRNA delivery using CED. **C)** Number of sgRNA positive cells for each sgRNA perturbation across cell types identified in (B). **D)** Integrated scRNA-seq UMAP of malignant and stromal/microenvironment cells from (C) overlayed with sgRNA UMI (left), KRAB expression level (middle), or whether the single cells were FACS sorted (right). **E)** Bubble plot of gene set enrichment analyses of gene expression modules (rows) following sgRNA perturbations (columns) spanning cell types identified in (C). Expression represents log2 fold change normalized to non-targeting controls from the same cell type. **F)** Differential communication probabilities for ligand-receptor interactions inferred from transcriptional profiles of perturbed macrophages (either negative control or *Ptprc* sgRNA) along with target cells of the tumor microenvironment within the same SB28 tumors.

Quantification of transcriptomic phenotypes for each CRISPRi genetic perturbation following CED in cell types with sufficient sgRNA coverage – macrophages, microglia, astrocytes, oligodendrocytes, oligodendrocyte progenitor cells (OPCs) – revealed cell type-dependent gene module alterations (Fig. 5E). We then investigated the consequences of perturbing the phosphatase *Ptprc* in macrophages *in vivo*, given its proinflammatory phenotype spanning multiple gene modules, which were not observed after perturbing this same target in other cell types (Fig. 5E). *Ptprc* knockdown resulted in upregulation of genes associated with interferon α and interferon γ response, as well as IL2/STAT5 and TNFα signaling (Fig. 5E), consistent with the role of *Ptprc* in innate immunity and cytokine signaling^50,51^. To determine if genetic perturbation could influence cell-cell interactions within SB28 intracranial tumors, we identified putative cell-cell interactions using CellChat^52^ (Methods) and quantified the pairwise differences in communication probabilities in macrophages following *Ptprc* suppression and other cell types, versus macrophages expressing non-targeting control sgRNAs and other cell types. Following *Ptprc* knockdown, *Ccl5* – *Ccr1*/*3*/*5* interactions increased with immune and stromal cell types (Fig. 5F), reflective of the overall proinflammatory phenotype of *Ptprc* knockdown in macrophages (Fig. 5E). In contrast, *Ptprc* knockdown in macrophages also abrogated cell-cell interactions through suppression of *Thbs1*, *Ccl6, Ccl9,* and *Ptprc* itself, reducing communication probabilities between macrophages and dendritic cells, microglia, SB28 malignant cells, among other cell types (Fig. 5F). These results not only reveal the role of *Ptprc* as a regulator of paracrine interactions within tumors, but also demonstrate that non-cancer cell types present in the tumor microenvironment can be perturbed and then phenotyped using our *in vivo* CED perturb-seq approach.

## Discussion

Multiplex *in vivo* perturbations coupled with measurement of rich transcriptional phenotypes can reveal genotype-phenotype relationships in normal and disease contexts that would not be feasible in cell cultures. Here, we developed a platform for *in vivo* perturb-seq to interrogate the function of oncogenic drivers and treatment response mechanisms in GBM, a fatal brain malignancy. Using iterative progression of *in vitro* target gene nomination, *in vivo* transplantation of *in vitro* infected malignant tumor cells, to direct delivery of sgRNA cargo using CED, we demonstrate that functional genomic screens performed *in vivo* reveal distinct phenotypic alterations spanning a diversity of cell autonomous and non-cell autonomous transcriptional pathways. *In vivo* CED perturb-seq also reveals how radiotherapy treatment of established tumors rewires transcriptional responses to genetic perturbations. Furthermore, this platform allows for interrogation of cellular interactions in the tumor microenvironment, highlighting the utility of perturbing cells that exist in their intact environment.

The observation that *in vivo* perturbations resulted in greater heterogeneity of transcriptional responses, especially when combined with radiotherapy, was a distinct feature of performing perturbations within a preformed tumor with an intact immune system. However, we cannot exclude contributions from differences in the experimental timelines between *in vitro* and *in vivo* perturb-seq workflows, as *in vivo* experimental timelines were constrained by slower growth rates of tumors compared to their respective cell cultures^53^, therefore requiring longer periods of time between perturbation and cell isolation. Furthermore, the apparent discrepancy between the cell types transduced by sgRNA lentiviruses in the GL261 CED experiments for CRISPRi in malignant cells (Fig. 4D, Fig. S4G) compared to the SB28 CED experiments for CRISPRi in the microenvironment cells (Fig. 5B, D) may be due to absence or presence of CRISPRi machinery, which is known to stabilize and substantially prolong the half-life of sgRNAs^54^. As such, the SB28 microenvironment experiments, which utilized animals that expressed H11-dCas9-KRAB in each normal cell, enabled recovery of sgRNAs in multiple glial and immune cell types (Fig. 5B, D). CRISPRi activity in the microenvironment cells also appeared less efficacious than in GBM malignant cells (Fig. S7B). This is likely a consequence of using computationally predicted sgRNAs for the microenvironment perturbations rather than sgRNAs nominated based on screen scores from genome-scale CRISPRi screens, which were feasible for cultured GBM cells (Fig. 1A, B). In future work, sgRNA nomination for perturb-seq could be performed entirely *in vivo*, thereby maximizing the likelihood of discovering *in vivo* specific phenotypes using highly active sgRNAs. Such efforts could be enabled by expanding or modifying the *in vivo* tropism of sgRNA libraries through adeno-associated viruses or virus-like particles^17,55^.

## Conclusions

In conclusion, *in vivo* CED perturb-seq, which we have iteratively developed, enables multiplex interrogation of complex biological processes in cancer and non-cancer cell types. This platform reveals mechanisms of treatment responses in GBM, and it also serves as the foundation for simultaneous discovery and characterization of therapeutic vulnerabilities that are uniquely functional *in vivo*.

## Methods

### In vitro CRISPRi screens, analysis, and validation

HEK-293T cells were cultured in Dulbecco’s Modified Eagle Medium (Gibco, #11960069) supplemented with 10% fetal bovine serum (FBS) (Life Technologies, #16141). Cell cultures were authenticated by STR analysis at the UC Berkeley DNA Sequencing Facility, as well as routinely tested for mycoplasma using the MycoAlert Detection Kit (Lonza, #75866-212). GL261 cells were cultured in Dulbecco’s Modified Eagle Medium (Gibco, #11960069) supplemented with 10% fetal bovine serum (FBS) (Life Technologies, #16141), and GL261 cells stably expressing the CRISPR interference (CRISPRi) machinery dCas9-KRAB were generated as previously described^56,57^.

Lentivirus was produced from transfected HEK293T cells with packaging vectors (pMD2.G #12259, Addgene, and pCMV-dR8.91, Trono Lab) following the manufacturers protocol (#MIR6605, Mirus). GL261 GBM cells were transduced with lentivirus harboring SFFV-dCas9-BFP-KRAB, and the top ∼25% of cells expressing BFP were FACS sorted, expanded, and FACS sorted a second time.

For genome-scale CRISPRi screening *in vitro*, we used a mouse sgRNA library^39^ comprising 27,300 sgRNAs targeting 5,234 cancer related and/or druggable genes, in addition to 530 non-targeting control sgRNAs. Pooled lentivirus was generated as above, and GL261 cells were transduced using spin-infection of viral supernatant of a MOI of ∼0.1 at 1000g for 120 minutes. 4 days of puromycin selection (1.5 µg/mL) was performed, followed by 2 days of growth in non-puromycin 10% FBS in DMEM media. Two replicates of each screen were performed at a coverage of 525x cells per sgRNA, in both no radiation and radiation (2 Gy x 5 fractions delivered every other day) treatment conditions. Infection efficiency was evaluated by measuring GFP positivity on flow cytometry. Initial (T0) cell populations were then frozen in 10% DMSO and processed for genomic DNA using the NucleoSpin Blood XL Kit (Machery-Nagel, #740950.50). Endpoint cell pellets were harvested for genomic DNA after 12 days of growth, corresponding to ∼7 and ∼5 doublings in the no radiation and radiation conditions, respectively. sgRNA sequencing libraries were prepared using NEBNext Ultra II Q5 PCR MasterMix (New England Biolabs, #M0544L) and sequenced on an Illumina NextSeq-500 as previously described^59^.

sgRNAs with fewer than 100 reads at T0 were removed from subsequent analysis. Enrichment or depletion of sgRNA abundances were determined by down sampling trimmed sequencing reads to equivalent amounts across all samples. Growth phenotype (gamma) was defined as log2(sgRNA count T12 (0Gy) / sgRNA count T0) minus median sgNTC log2(sgRNA count T12 (0Gy) / sgRNA count T0), then normalized by the number of cell doublings, as previously described^56^. Radiation phenotype (tau) was defined as log2(sgRNA count T12 (2Gy x 5) / sgRNA count T0) minus median sgNTC log2(sgRNA count T12 (2Gy x 5) / sgRNA count T0), then normalized by the number of cell doublings in the radiation screen. Radiation:growth ratio phenotype (rho) was defined as log2(sgRNA count T12 (2Gy x 5) / sgRNA count T12 (0 Gy)). Gene-level phenotypes were summarized as the mean of the top 3 sgRNAs against a given gene, ranked according to screen phenotype. Statistical significance was calculated using the Mann-Whitney-U test for a given perturbation compared to the sgRNA distribution of the non-targeting control sgRNAs (Table S1). A discriminant threshold of 7, derived from the product of normalized gene phenotype and -log10(p-value), corresponding to an empiric false discovery rate of ∼2%, was selected for hit definitions^56^. To ascertain the fidelity of our mouse cell CRISPRi screens, we overlapped gamma phenotypes (without radiotherapy treatment) in our GL261 screens with gamma phenotypes from K562 CRISPRi cells subjected to genome-wide human CRISPRi growth screens^56^.

Internally controlled competitive growth assays were then performed to validate CRISPRi screen hits. GL261 cultures were partially transduced with sgRNA expression lentiviruses with a GFP tag (Addgene 187241)^40^, and the percentage of sgRNA positive cells were measured over time using flow cytometry, in the presence or absence of radiotherapy (2 Gy x 5 fractions).

### Intracerebral tumor establishment

Five to six-week-old female C57/B6 (Envigo Laboratories, Livermore, CA), housed under aseptic conditions, received intracranial tumor cell injection as previously described^58^, and as approved by the University of California San Francisco Institutional Animal Care and Use Committee. For the microenvironment perturb-seq, mice expressing dCas9-KRAB from the H11 locus with a mCherry tag (JAX # 030000) were utilized. Briefly, mice were anesthetized by combination of intraperitoneal injection of a mixture containing ketamine (100 mg/kg) and xylazine (10 mg/kg), and inhalation of isoflurane. The scalp was surgically prepped, and a skin incision ∼10 mm in length was made over the middle frontal to parietal bone. The surface of the skull was exposed so that a small hole was made 3.0mm to the right of the bregma and just in front of the coronal suture with a 25-gauge needle. A 26-gauge needle attached to a Hamilton syringe was inserted into the hole in the skull. The needle was covered with a sleeve that limits the injection depth to 3-4mm. 3 µL of tumor cell suspension (300,000 GL261 cells; 30,000 SB28 cells) was injected into the right caudate putamen at a rate of 1 µL/min by free hand. The skull surface was then swabbed with hydrogen peroxide before the hole was sealed with bone wax to prevent reflux. The scalp was closed with surgical staples.

### Bioluminescence imaging of intracranial tumor growth

For bioluminescence imaging (BLI), mice were anesthetized with inhalation of isoflurane, then administered 150 mg/kg of luciferin (D-luciferin potassium salt, Gold Biotechnology, St. Louis, MO) via intraperitoneal injection. Ten minutes after luciferin injection, mice were examined for tumor bioluminescence with an IVIS Lumina imaging station and Living Image software (Caliper Life Sciences, Alameda, CA), and intracranial regions of interest were recorded as photons per second per steradian per square cm^58^.

### Convection enhanced delivery of lentivirus

Convection enhanced delivery was performed as previously described^59^. Briefly, infusion cannulas were constructed with silica tubing (Polymicro Technologies, Phoenix, AZ) fused to a 0.1 ml syringe (Plastic One, Roanoke, VA) with a 0.5-mm stepped tip needle that protruded from the silica guide base. The syringe was loaded with 15 µL concentrated lentivirus produced using the LV-MAX Lentiviral Production System (Thermo Fisher #A35684) according to manufacturer’s protocol, followed by 200x concentration using ultracentrifuge (Beckman L8-80M) at 25,000g for 2.5 hours at 4°C. Concentrated lentivirus was tested for *in vitro* titers and achieved 1-2 x 10^9^ TU/mL. The syringe was attached to a microinfusion pump (Bioanalytical Systems, Lafayette, Ind.), and the syringe with silica cannula was lowered through a puncture hole made in the skull^58,60^, targeting the same region in the caudate putamen where tumor cells had been previously injected. The concentrated lentivirus was infused at a rate of 1 µL/min until a volume of 15 µL had been delivered. Cannulas were removed 2 minutes after completion of infusion.

### Whole brain animal irradiation

Mice were anesthetized by combination of intraperitoneal injection of a mixture containing ketamine (100 mg/kg) and xylazine (10 mg/kg), and inhalation of 2.5% isoflurane with 1 liter of oxygen per minute for 5 minutes prior to being positioned on an irradiation platform located 16.3cm from a Cesium-137 source (J. L. Shepherd & Associates, San Fernando, CA). Animal subjects’ eyes, respiratory tracts, and bodies were protected with lead shielding. Whole brain irradiation (2 Gy for 5 daily fractions) was delivered at a dose rate of 247 cGy/min^61^. After irradiation, animals were monitored until recovery from anesthesia. Following no radiation or radiation treatment, animals were sacrificed and brains were quickly removed from the skull, and the tumor mass was microdissected as previously described^58^.

### Imaging

Fluorescence microscopy was performed on a Zeiss LSM 800 confocal laser scanning microscope with Airyscan. Images were processed and quantified from at least 2 regions per condition using ImageJ9. Hematoxylin and eosin stained slides were imaged on a Zeiss Axio Zoom V16 light microscope.

### In vitro and in vivo perturb-seq in malignant cells

Perturb-seq sgRNA libraries were designed by nominating 48 genes with radiation sensitizing, radiation resistance, negative growth, or positive growth phenotypes in *in vitro* CRISPRi screens as described above. Protospacer sequences were selected from the optimized mouse CRISPRi v2 library^39^, and the top two scoring sgRNAs for each target gene based on depletion or enrichment phenotypes were cloned into dual sgRNA lentivirus expression vectors with direct capture tags (Addgene 187241)^40^ using NEBuilder HiFi DNA Assembly Master Mix (New England Biolabs #E2621L) (Table S2). Concentrated lentivirus of pooled sgRNA libraries were produced as described above.

For *in vitro* perturb-seq, GL261 cells expressing dCas9-KRAB were transduced with lentivirus at an MOI of ∼0.1, and cells were FACS sorted (BD FACSAria Fusion) for GFP positivity 48 hours following transduction. Cell cultures were irradiated using a Cesium-137 source to 2 Gy x 5 fractions delivered daily, starting on same day of FACS. 12 hours following the final fraction of radiotherapy, cells were trypsinized and harvested in single cell suspension on the 10x Chromium Controller (10x Genomics, #1000204). Single-cell perturb-seq libraries were processed using the Chromium Next GEM Single Cell 3’ GEM, Library & Gel Bead Kit v3.1 with Feature Barcoding (10x Genomics, #1000269), allowing direct capture of modified sgRNAs, and sequenced on a illumina NovaSeq-6000. *In vitro* perturb-seq experiments were performed in biological duplicate cultures for each treatment condition.

For pre-infected *in vivo* perturb-seq, GL261 cells expressing dCas9-KRAB were transduced with lentivirus at an MOI of ∼0.1 and were subjected to puromycin selection (1.5 µg/uL) for 3 days, starting 2 days after transduction, followed by 2 days of growth in non-puromycin recovery media. 300,000 cells were intracranially injected into each mouse as described above. Tumors were allowed to establish and expand for 5 days, and radiation (or no treatment) was delivered to a dose of 2 Gy x 5 daily fractions as described above. For tumors of the no radiation arm, tumor harvest was performed 12 days following intracranial injection of cells. For tumors of the radiation arm, tumor harvest was performed 2 days following completion of radiation treatment, or 14 days total after intracranial injection of cells. Harvested tumors were minced and dissociated to single cell suspension using the Papain Dissociation System (Worthington #LK003150) following manufacturer’s protocol without the use of ovomucoid protease inhibitor. Cell suspensions were passed through a 70 μm strainer (Corning, #352350), centrifuged at 300g for 5 minutes, and resuspended in cold phosphate buffered saline. To capture cells of the tumor microenvironment as well as sgRNA transduced malignant cells *in vivo*, we processed a fraction of the dissociated tumor for immune cell depletion using CD11b MicroBeads (Miltenyi Biotec #130-097-142) following manufacturer’s protocol on LS Columns, followed by FACS sorting for sgRNA positive cells tagged with GFP. Sorted cells as well as a separate fraction of the dissociated tumor that was not CD11b depleted or FACS sorted were processed for scRNA-seq with direct capture of sgRNA tags using 10x Chromium Controller (10x Genomics, #1000204). Single cell perturb-seq libraries were processed as described for *in vitro* perturb-seq. Pre-infected *in vivo* perturb-seq experiments were performed in biological triplicate tumors.

For CED *in vivo* perturb-seq, GL261 cells expressing dCas9-KRAB were intracranially injected into mice as described above. Tumors were allowed to establish and expand for 5 days, and CED of concentrated lentivirus was performed as described above. For the radiation treatment arm, whole brain irradiation was delivered to a dose of 2 Gy x 5 daily fractions as described above, initiating 2 days following CED to allow lentiviral transduction and sgRNA expression. For tumors of the no radiation arm, tumor harvest was performed 5 days following CED, for a total of 12 days following implantation of cells. For tumors of the radiation arm, tumor harvest was performed 2 days following completion of radiation treatment, or 14 days total after intracranial injection of cells. Harvested tumors were minced and dissociated to single cell suspension as described for pre-infected *in vivo* perturb-seq. 5 tumors from biological replicate animals in each treatment arm were pooled to increase sgRNA positive cell recovery (0.7-1.1% sgRNA positive per pool by FACS). From these pools of dissociated cells, separate immune cell depletion followed by FACS sorting for GFP and unenriched populations were captured for scRNA-seq with direct capture of sgRNAs as described above. Single cell perturb-seq libraries were processed as described for *in vitro* perturb-seq.

### In vivo perturb-seq in tumor microenvironment cells

Microenvironment perturb-seq sgRNA libraries were designed by nominating 6 genes that were either overexpressed in myeloid cell types from scRNA-seq of SB28 orthotopic tumors, or by their known gene function in the myeloid lineage. Protospacer sequences were selected from the optimized mouse CRISPRi v2 library^39^ by their predicted rank order, and the top 3 sgRNAs were individually cloned into single sgRNA expression vectors with direct capture cs1 sequences (Addgene # 122238)^10^ using restriction digest (BstXI and BlpI) and T4 ligation (NEB # M0202M). A non-targeting sgRNA sequence was included as well. Concentrated lentivirus of pooled sgRNA was produced as described above. SB28 cells^45^ without CRISPRi machinery were intracranially injected into the H11-dCas9-KRAB mice (JAX # 030000) as described above, and CED of concentrated lentivirus was performed as described above 7 days after tumor implantation. 5 days following CED, tumors were harvested, minced, and dissociated to single cell suspension using the Papain Dissociation System (Worthington #LK003150) following manufacturer’s protocol without the use of ovomucoid protease inhibitor. Cell suspensions were passed through a 70 μm strainer (Corning, #352350), centrifuged at 300g for 5 minutes, and resuspended in cold phosphate buffered saline. To capture the full spectrum of cell types including sgRNA positive and negative cells, both single cell suspensions sorted for sgRNA positivity and lacking the SB28 GFP marker (RFP+/GFP-), as well as unsorted cells were processed for scRNA-seq with direct capture of sgRNA tags using 10x Chromium Controller (10x Genomics, #1000204). Single-cell perturb-seq libraries were processed using the Chromium Next GEM Single Cell 3’ GEM, Library & Gel Bead Kit v3.1 with Feature Barcoding (10x Genomics, #1000269), allowing direct capture of modified sgRNAs, and sequenced on a illumina NovaSeq-6000. Tumor microenvironment perturb-seq was performed in biological quadruplicates, with each replicate consisting of two pooled animals.

## Perturb-seq computational analyses

### Pre-processing, sgRNA calling, cell type identification

Library demultiplexing, gene expression read alignment to human genome GRCh38, UMI quantification, and sgRNA assignment and quantification were performed in Cell Ranger version 6.1.2 with sgRNA barcoding (10X Genomics). Single cell RNA-seq analysis was performed in Seurat version 4.3.0^62^ in R version 4.3.1. For analysis of cellular heterogeneity across all perturb-seq experiments, including malignant cells and microenvironment, cells with greater than 200 detected features were retained. RNA expression data was transformed using SCTransform^63^ in Seurat, and expression featureplots show log2 UMI’s corrected by SCTransform. Expression heatmaps of scRNA-seq data show SCTransform residuals (normalized expression). Uniform manifold approximation and projection (UMAP) was performed using the Seurat function RunUMAP using the first 30 dimensions from principal component analysis of the transformed expression data, using parameters (min.dist = 0.7). Cell type identification of stromal and microenvironment cells was performed using single cell multiresolution marker-based annotation (scMRMA) version 1.0^64^, using the PanglaoDB mouse cell types reference^65^, with manual confirmation of known marker gene specificity. To deconvolve malignant tumor cells from stromal tissue with potentially similar RNA expression profiles, we integrated scRNA-seq data from our *in vitro* cultures and *in vivo* experiments to allow manual annotation of *in vitro* GBM cells, as these were processed in separate 10x lanes and therefore could be distinguished by metadata. Similarly, sgRNA positive cells from the pre-infected perturb-seq experiments could be readily identified as malignant GBM cells, as these were transduced with sgRNA prior to *in vivo* implantation. Using these pre-defined cell clusters of malignant cells, projection of cells from the CED perturb-seq experiment onto an integrated UMAP space demonstrated a highly abundant population of cells which co-clustered with malignant cells from the pre-infected perturb-seq experiment, and therefore these cells were identified as *in vivo* malignant GBM cells and were therefore analyzed in the context of their corresponding non-malignant cells.

Cells were retained for further analysis if they expressed both the expected sgRNA A and sgRNA B against a given target gene, corresponding to the sgRNAs of the dual sgRNA expression vector. Target gene knockdown was quantified by library normalizing the untransformed transcriptome UMIs of each sgRNA positive cell and obtaining the mean expression of each gene across all cells belonging to a given sgRNA within each individual GEM group. RNA remaining for each gene target was calculated by dividing the pseudobulk expression in on-target cells with cells expressing non-targeting negative control sgRNAs, with a pseudocount of 0.1 added to each component. Quantification of RNA remaining values were capped at a ceiling of 1.0. Expression featureplots were generated using SCPubr version 1.1.2^66^.

### Differential gene expression analysis

First, we filtered the expression objects as follows: For each differential expression run, we isolated cells by the desired experimental context (*in vitro,* pre-infected, CED), selected for FACS sorted only cells in the pre-infected context, and selected cells with sgRNA expression called for both expected sgRNAs against a given target gene. We removed cells with perturbations that had coverage of less than or equal to 5 cells. Next, we used the default findDE method from the DElegate package version 1.1.0^67^ to find differentially expressed genes. DElegate is a wrapper around DESeq2 that adapts bulk sequencing methods for single cell data. The effect of running findDE is to run DESeq2 with the Wald test on randomly assigned 3-group pseudoreplicates. findDE’s input is a Seurat object and metadata specifications for a group of interest and a control group to measure against. We ran findDE comparing cells with a particular perturbation +/-radiation to the non-targeting perturbation + no radiation—a normalization scheme we call “noRTNormalized”—as well as a particular perturbation +/-radiation to the treatment-matched non-targeting perturbation +/-radiation, a normalization scheme that internally controls for whether cells received radiotherapy, we call “condNormalized.”. Per perturbation, findDE’s output is a table that provides—for each gene expressed in the cells compared—the following information: the gene name, the average expression of the gene (using deseq:baseMean), the log2 fold change, the test statistic, the p-value, and the FDR adjusted p-value. We performed this DElegate::findDE workflow for all perturbations across the three experimental contexts—*in vitro,* pre-infected, and convection enhanced delivery. Each output represented a comparison of expression between perturbed cells and the control cells. Next, we screened each of the resulting results sets to find the differentially expressed genes. Within each set, we considered a gene differentially expressed if it passed DESeq2’s expression filters (meaning it had a non-NA p-value and FDR adjusted p-value), had an absolute value log2 fold change of greater than 0.1, and an adjusted p-value of less than 0.05.

### Downsampled differential gene expression analysis

To account for differences in cell sampling between radiotherapy and no radiotherapy treatment conditions, we prepared downsampled differential expression output by modifying the preprocessing steps prior to running DElegate’s findDE function. First, we removed perturbations with less than or equal to 5 cells. Then we randomly downsampled using dplyr::sample_n the number of cells for each perturbation such that there were equal numbers of cells in the radiotherapy and no radiotherapy conditions, for each experimental context (ie. *in vitro* vs. *in vivo*). Differential gene expression analysis was then performed as described above, and the number of differentially expressed genes in the downsampled analysis was quantified identically to the non-downsampled workflow.

### Gene Set Enrichment Analysis (GSEA)

For GSEA, we used the DESeq2 output as described above. For each perturbation x radiation condition—and thus each comparison or each CSV—we first used the annotables package (v0.2.0) to convert gene symbols to ENSEMBL v109 gene IDs. Genes that failed to convert were eliminated. We also removed genes that DESeq2 indicated didn’t pass independent filtering. Then we ranked the remaining genes by their log2 fold changes and ran gene set enrichment analysis using fgsea (v1.27.1), a package for fast GSEA, and msigdbr (v7.5.1), the R implementation of the Molecular Signatures Database. We set the maximum size of allowed gene sets to 500 and set the eps parameter—which bounds the lowest p-value possible—to 0 to allow for arbitrarily low p-values.

This analysis resulted in a set of pathways for each perturbation x radiation condition, their corresponding normalized enrichment scores, and the p-values and adjusted p-values indicating significance of enrichment. We ran this analysis for both “noRTNormalized” and “condNormalized” schemes, described above. We indicated the normalized enrichment scores for all pathways that have an adjusted p-value lower than 0.05 in any perturbation x radiation condition.

For bubble plots, we show the mean log2 fold change, which we calculated as follows for each pathway x (perturbation x radiation) condition: first, we found the genes shared between a given pathway’s msigdb gene set and the union of all differentially expressed genes with valid ENSEMBL IDs across perturbation x radiation conditions for a given normalization scheme. Then, we took the mean log2 fold change of these shared genes for each perturbation x radiation condition. Genes that did not pass DESeq quality filtering were set to have a log2 fold change of 0 prior to calculating the mean, to ensure that genes that didn’t pass quality filtering would have minimal contribution to the mean log2 fold changes.

### Visualization of perturbation space with linear discriminant analysis

To separate perturbations in transcriptomic space, we ran linear discriminant analysis (LDA) on all perturbations for each radiotherapy/no radiotherapy condition. We first filtered out low coverage perturbations (<= 5 cells) and applied a centered log transformation normalization. Then, we used Seurat’s^68^ CalcPerturbSig function to generate perturbation signatures based on 40 principal components and the 20 nearest neighbors. Finally, we separated out cells by treatment condition and fed them into Seurat’s MixscapeLDA function. We used a log2 fold change threshold of 0.1 to be consistent with our DESeq2 settings. Otherwise, we set the seed to the default value of 42 and the number of principal components to the default value of 10. For visualization, we chose a UMAP with the maximum number of linear discriminants found per condition. To perform LDA UMAP analysis with the downsampled set of cells, we first removed perturbations with less than or equal to 5 cells. Then we randomly downsampled the number of cells for each perturbation using the R function dplyr::sample_n such that there were equal numbers of cells in the radiotherapy and no radiotherapy conditions for each experimental context. We then performed linear discriminant analysis with the same parameters as above.

To quantify density of cells in UMAP space, we calculated gaussian kernel densities for the UMAP coordinates in 2 dimensions for all (or downsampled) cells expressing a given sgRNA pair, with 10,000 total bins (100 bins in each UMAP 1/UMAP 2 dimension spanning the full range of coordinates). The maximum z density was obtained for each sgRNA target in either radiotherapy or no radiotherapy conditions, and the difference between these values was calculated.

### Treatment-perturbation modeling and synergy analysis

To model interactions between perturbations and radiation treatment, we used CINEMA-OT (causal independent effect module attribution with optimal transport), a method that separates treatment and confounder effects using independent component analysis, then matches cells across treated/untreated conditions using optimal transport^43^. The matching can then be used to generate individual treatment effect matrices (ITE matrices), which indicate what a hypothetical treatment effect would be for each gene of each control cell. When multiple treatments are present, one can use the matchings to quantify a synergy score between treatments by defining a synergy matrix as ITE(A + B) -(ITE(A) + ITE(B)), where A and B are separate treatments. We used cinemaot.synergy in the cinemaot (v0.0.5) Python package (https://github.com/vandijklab/CINEMA-OT), to generate synergy scores per cell per gene between sgRNA perturbations and radiation. This was performed by passing in a perturbation with radiation as A + B, the same perturbation with no radiation as A, and radiation with a non-targeting sgRNA as B. Before running CINEMA-OT, we generated highly variable genes using scanpy’s (v1.9.6) highly_variable_genes method, choosing a min_mean of 0.0125, max_mean of 3, min_disp of 0.5, max_disp of Inf, span of 0.3, and 20 bins. We then ran PCA on the highly variable genes using scanpy’s^68^ tl.pca method, passing in the “arpack” solver, 50 dimensions, a random state of 0, and “zero-centered,” meaning PCA calculated from the covariance matrix.

CINEMA-OT itself—and thus the cinemaot.synergy method—takes in the following parameters: the number of independent components, a threshold for setting the Chatterjee coefficient for confounder separation, a parameter for setting the smoothness of entropy-regularized optimal transport, a parameter for the stop condition of OT convergence, and whether to return matrices weighted by the number of cells (a method termed CINEMA-OT-W). We set the number of independent components to 10, the threshold for confounder separation to 0.5, and the smoothness parameter to 1e-3 to adjust for data sparsity. The stop condition we maintained at the default value of 1e-3, and we did not weight by the number of cells.

We ran cinemaot.synergy for each perturbation across all experimental contexts, generating a synergy score per cell per gene—a synergy score matrix. We then calculated synergy scores per cell by summing the synergies for any gene whose absolute value (average synergy) across all cells in that perturbation was greater than the default parameter of 0.15. For visualization purposes, we then calculated the L2-norms for each cell.

### Microenvironment perturb-seq analysis

Preprocessing, data transformation, and UMAP analysis was performed using Cell Ranger version 6.1.2 with sgRNA barcoding (10X Genomics) followed by Seurat version 4.3.0^62^ in R version 4.3.1 as described above. For microenvironment cells, cells with greater than 200 detected features were retained. Cell types were identified by performing Louvain clustering using the first 30 dimensions from PCA space and using a resolution parameter of 0.4. Top marker genes from these clusters were queried in EnrichR^70,71^. Unbiased cell type annotation was also performed using scMRMA as described above, with good agreement to cluster-based annotations. SB28 cells were readily identified by overexpression of *Nras*^45^. On target gene knockdown analysis was performed as described above.

For differential gene expression analysis in the microenvironment cells, instead of running DESeq independently for each radiation condition, we ran DESeq independently for each cell type. We first filtered the expression objects, removing cells with perturbations with coverage less than or equal to 5 cells or cells that do not have expression of a single sgRNA. We then ran DElegate’s findDE method for each perturbation in each cell type against the non-targeting sgRNA (sgNegCtrl3). Per perturbation, we again received a CSV file that provided gene names, log2 fold changes, adjusted p-values, etc. Each CSV file was fed into downstream analysis. Gene set enrichment analysis was performed as described above. However, instead of running GSEA independently per radiation condition, we ran GSEA independently for each cell type.

Cell-cell interaction analysis between ligands and receptors was performed using CellChat version 1.6.1^52^, using the CellChatDB.mouse interaction database. Overexpressed genes and interactions were identified, and communication probability was determined using a truncatedMean approach with trim = 0.01, a minimum of 5 cells per communication, without weighting of relative population size. Interactions with p value < 0.05 were retained for analysis. To calculate differential communication probabilities following a given sgRNA perturbation, we generated matrices corresponding to the difference in communication probability between perturbed and non-targeting control cells for each ligand-receptor combination across all cell types. For heatmap visualization, only ligand-receptor combinations with differential communication probabilities greater than 0.05 in magnitude in at least one cell type were retained.

---

**Table S1. *In vitro* growth and radiation phenotypes from pooled CRISPRi screens in GL261**

**Table S2. *sgRNA libraries used for in vivo perturb-seq***

## Declarations

### Ethics approval and consent to participate

Study was approved by the University of California San Francisco Institutional Animal Care and Use Committee.

### Consent for publication

Not applicable.

### Availability of data and materials

All sequencing data have been deposited on NCBI SRA accession number PRJNA1005229. Computational code has been deposited on Github at https://github.com/GilbertLabUCSF/gbm_perturb.

### Authors’ contributions

All authors contributed substantially to the inception, design, implementation, analysis, or writing of this report. All authors approved the manuscript. SJL, LG, DRR, WW, and TO designed the study and analyses. Experiments were performed by SJL, JP, TCC, KS, TO’L, TO, DRR, and LG. Computational data analyses were performed by SJL, CZ, AB, and DW. The study was supervised by SJL, DL, TO, MSB, WW, DRR, and LG. The manuscript was prepared by SJL, LG, and CZ with input from all authors.

## Acknowledgements

As described above.

---

**Figure S1.**
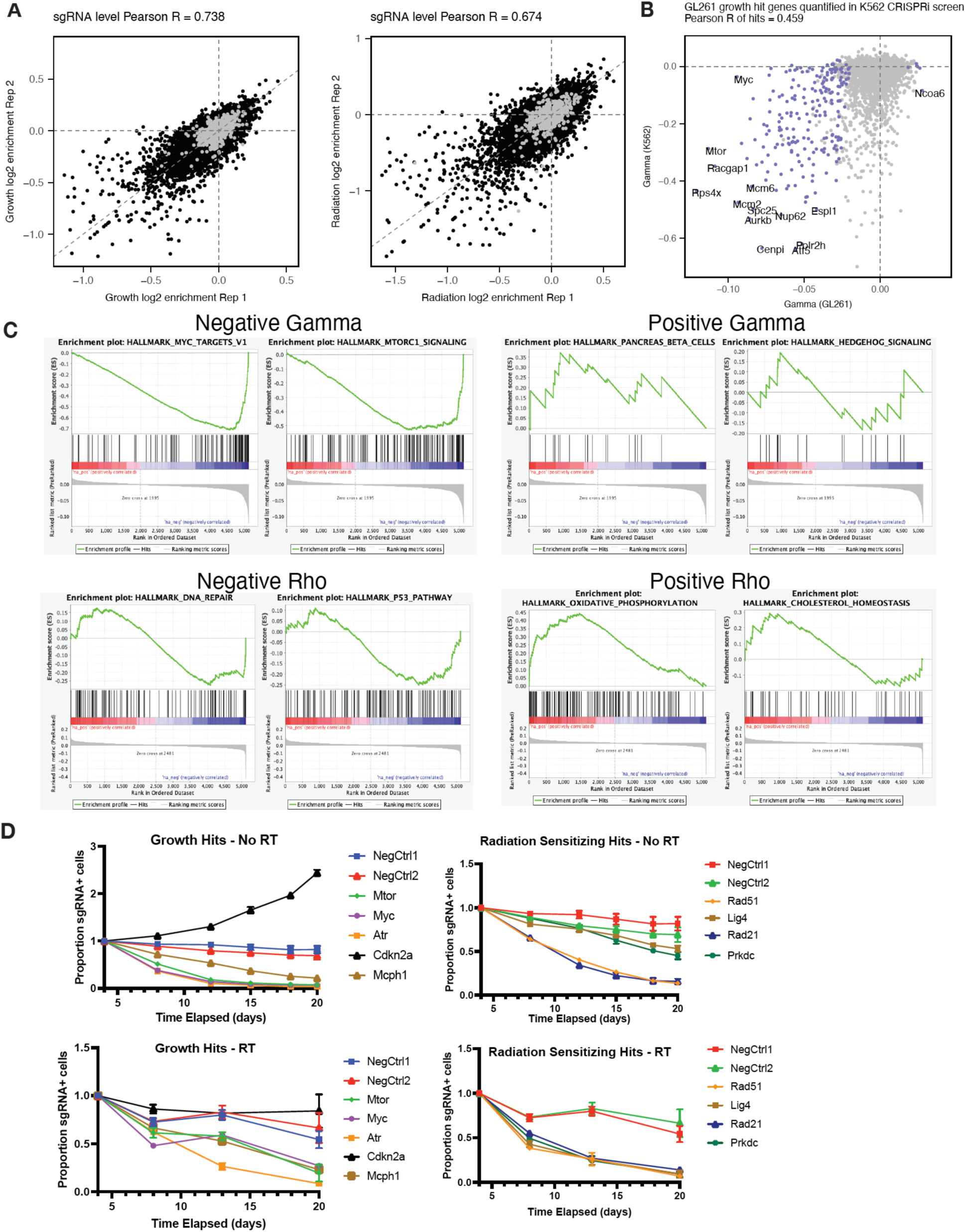
CRISPRi screen reproducibility and validation in GL261 GBM cells. **A)** Inter-replicate scatter plot of growth (left) and radiation (right) phenotypes for sgRNAs targeting genes (black) and non-targeting negative control sgRNAs (gray). **B)** Comparison of normalized growth phenotypes (gamma) in GL261 cells compared to published growth phenotypes in K562 cells from Horlbeck et al. 2016. **C)** Gene set enrichment analyses (GSEA) for negative (left) or positive (right) gamma (top) or rho (bottom) hit genes. **D)** Internally controlled competitive growth assays validating growth (left) and radiation sensitizing (right) hit genes in the absence (top) or presence (bottom) of fractionated radiotherapy (2 Gy x 5 fractions) in GL261 cells expressing CRISPRi machinery.

**Figure S2.**
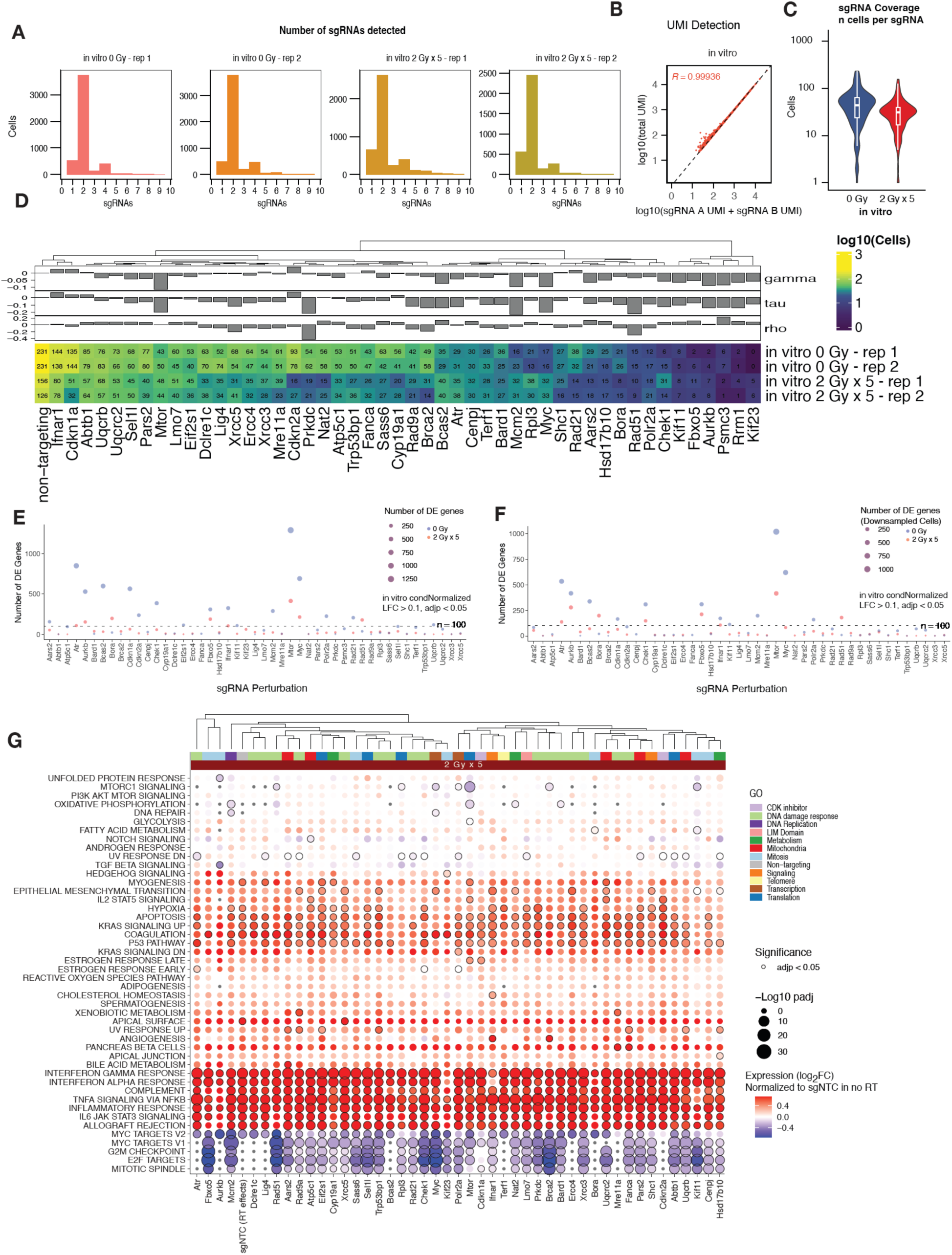
Quality control metrics for *in vitro* perturb-seq. **A)** Histogram of sgRNA detection in single cells positive for sgRNA expression following transduction with dual sgRNA lentiviruses. **B)** Scatter plot comparing total sgRNA UMI’s to the sum of UMI’s corresponding to the expected sgRNA A and sgRNA B for each targeted gene, for *in vitro* perturb-seq experiments. **C)** Distribution of the number of cells expressing each sgRNA among GL261 malignant cells from *in vitro* perturb-seq. **D)** Heatmap of per-sgRNA coverage for each targeted gene in GL261 *in vitro* perturb-seq. Numbers represent number of cells expressing both sgRNAs against each target gene. Top bar charts show growth (gamma), radiation (tau), and radiation:growth ratio (rho) *in vitro* screen phenotypes. Each row of the heatmap shows an individual replicate experiment for the *in vitro* experiment (n = 2 biological replicate cultures per condition). **E)** Number of differentially expressed genes for each perturbation in no radiotherapy (blue) or radiotherapy (2 Gy x 5 fractions; red) conditions. **F)** As in (E) after downsampling cells for differential expression analysis such that radiotherapy and no radiotherapy conditions had equivalent numbers of cells for each sgRNA target. **G**) Bubble plot of gene set enrichment analyses of gene expression modules (rows) following sgRNA perturbations (columns) with radiotherapy. Expression represents log2 fold change normalized to non-targeting controls from the no radiotherapy condition.

**Figure S3.**
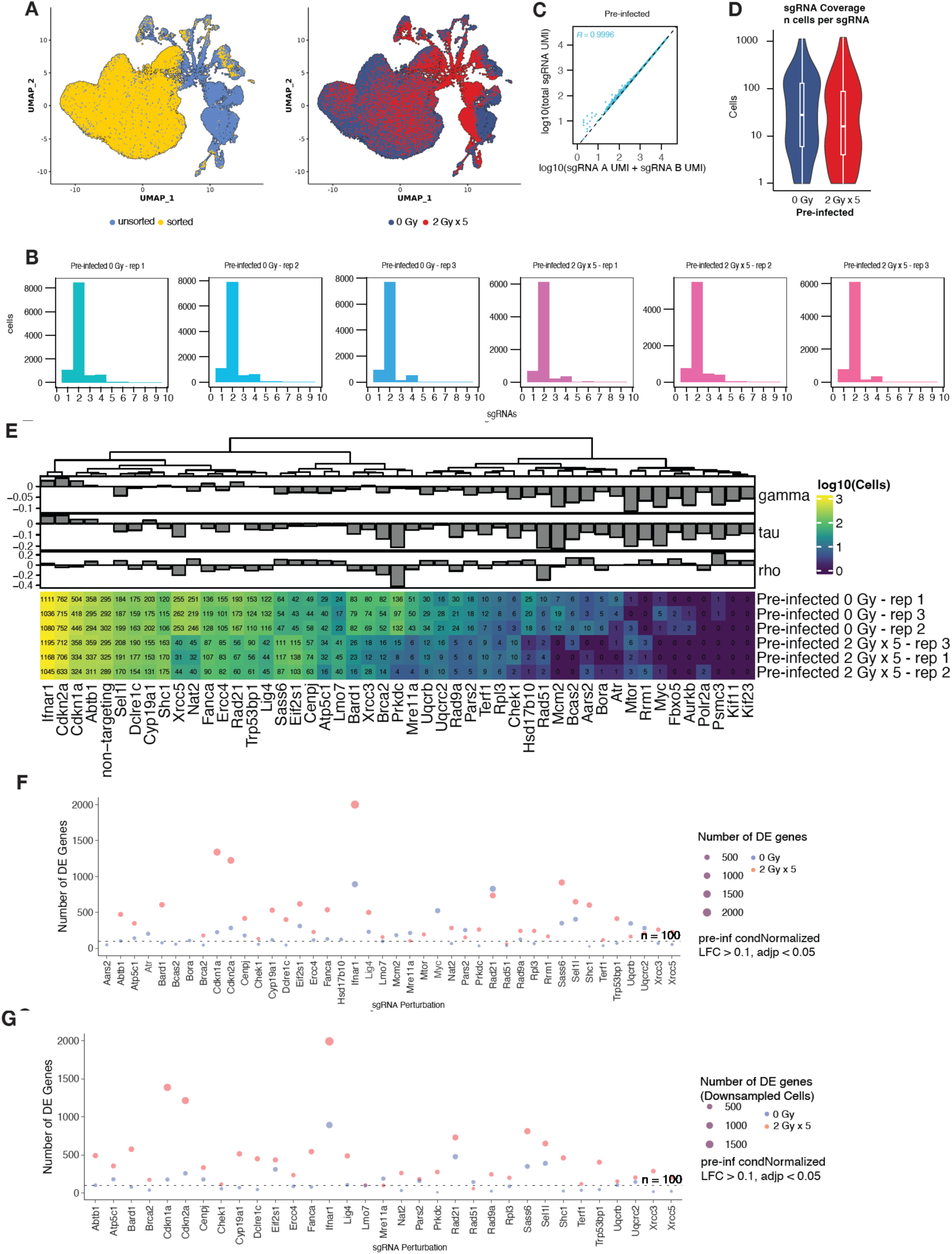
Quality control metrics for *in vivo* perturb-seq using pre-infected cells. A) Integrated scRNA-seq UMAP of malignant and stromal/microenvironment cells from *in vivo* pre-infected perturb-seq experiments colored by whether the single cells were negative MACS selected against Cd11b cells and then FACS sorted for sgRNA expression (left) or by radiotherapy condition (right). B) Histogram of sgRNA detection in single cells positive for sgRNA expression following transduction with dual sgRNA lentiviruses and transplanted intracranially to form orthotopic allografts. C) Scatter plot comparing total sgRNA UMI’s to the sum of UMI’s corresponding to the expected sgRNA A and sgRNA B for each targeted gene, for *in vivo* pre-infected perturb-seq experiments. D) Distribution of the number of cells expressing each sgRNA among GL261 malignant cells from *in vivo* pre-infected perturb-seq. E) Heatmap of per-sgRNA coverage for each targeted gene in GL261 *in vivo* pre-infected perturb-seq. Numbers represent number of cells expressing both sgRNAs against each target gene. Top bar charts show growth (gamma), radiation (tau), and radiation:growth ratio (rho) *in vitro* screen phenotypes. Each row of the heatmap shows an individual replicate experiment for the *in vivo* pre-infected experiment (n = 3 biological replicates per condition). F) Number of differentially expressed genes for each perturbation in no radiotherapy (blue) or radiotherapy (2 Gy x 5 fractions; red) conditions. G) As in (F) after downsampling cells for differential expression analysis such that radiotherapy and no radiotherapy conditions had equivalent numbers of cells for each sgRNA.

**Figure S4.**
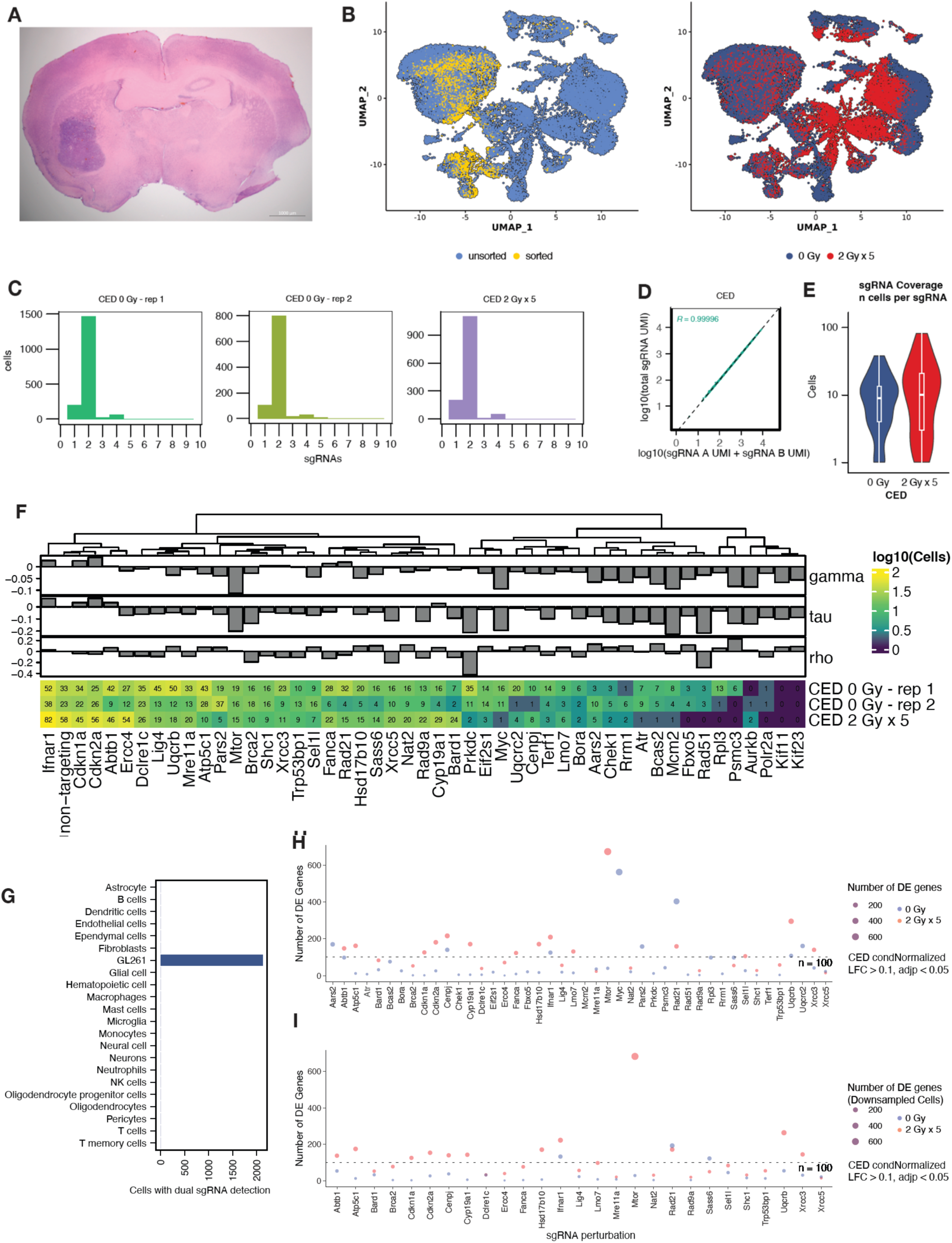
Quality control metrics for *in vivo* perturb-seq using convection enhanced **delivery. A)** Hematoxylin and eosin staining of GL261 orthotopic tumor centered in the right striatum. Scale bar, 1000 μm. **B)** Integrated scRNA-seq UMAP of malignant and stromal/microenvironment cells from *in vivo* CED perturb-seq experiments colored by whether the single cells were negative MACS selected against Cd11b cells and then FACS sorted for sgRNA expression (left) or by radiotherapy condition (right). **C)** Histogram of sgRNA detection in single cells positive for sgRNA expression following CED with dual sgRNA lentiviruses directed into orthotopic allografts. **D)** Scatter plots comparing total sgRNA UMI’s to the sum of UMI’s corresponding to the expected sgRNA A and sgRNA B for each targeted gene, for *in vivo* CED perturb-seq experiments. **E)** Distribution of the number of cells expressing each sgRNA among GL261 malignant cells from *in vivo* CED perturb-seq. **F)** Heatmap of per-sgRNA coverage for each targeted gene in GL261 *in vivo* CED perturb-seq. Numbers represent number of cells expressing both sgRNAs against each target gene. Top bar charts show growth (gamma), radiation (tau), and radiation:growth ratio (rho) *in vitro* screen phenotypes. Each row of the heatmap shows an individual replicate experiment for the *in vivo* CED experiment (n = 5 animals pooled per replicate spanning both conditions). **G)** Number of cells pertaining to each cell type from GL261 tumors with successful dual sgRNA recovery. **H)** Number of differentially expressed genes for each perturbation in no radiotherapy (blue) or radiotherapy (2 Gy x 5 fractions; red) conditions. **I)** As in (H) after downsampling cells for differential expression analysis such that radiotherapy and no radiotherapy conditions had equivalent numbers of cells for each sgRNA.

**Figure S5.**
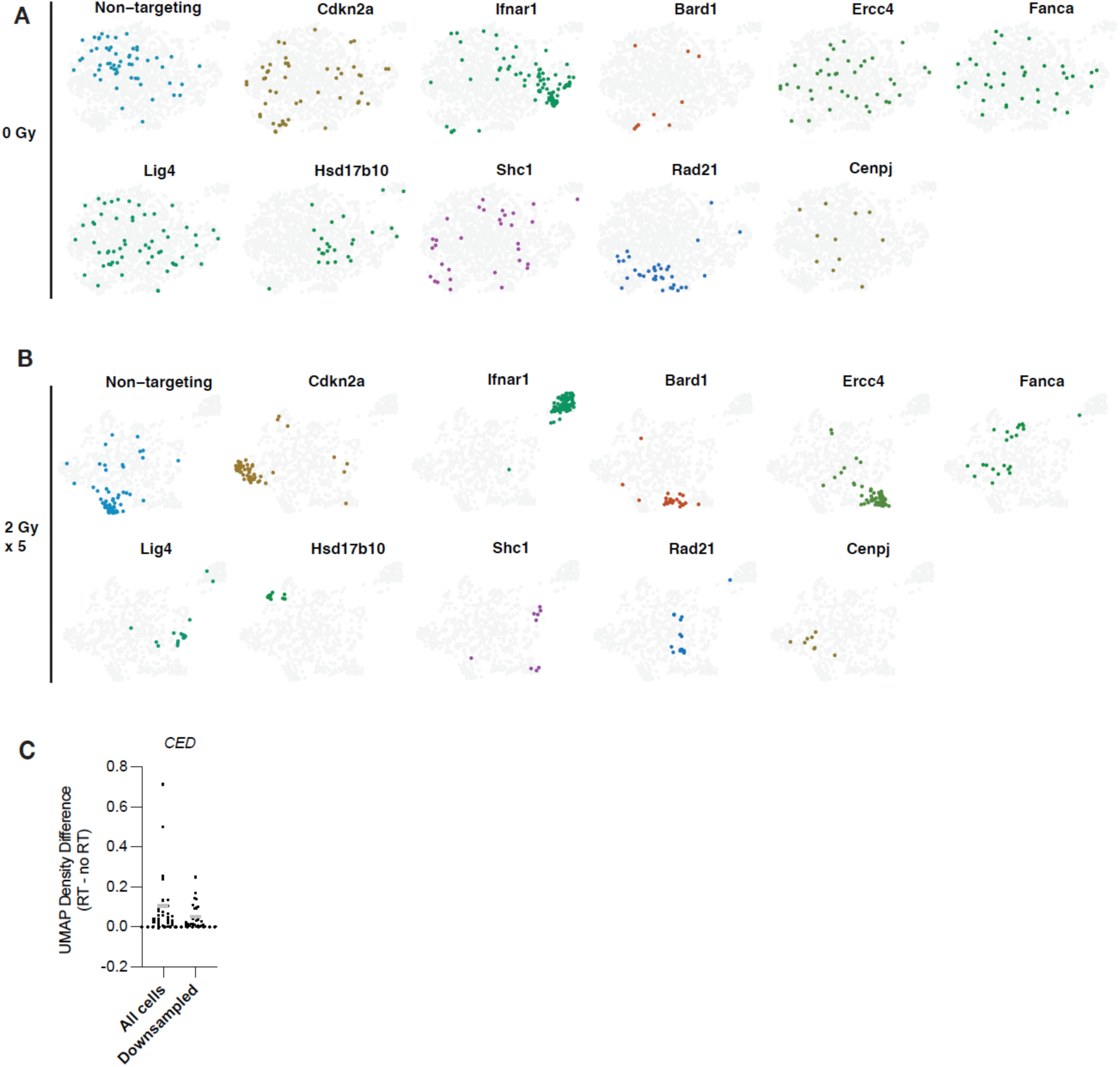
Linear discriminant analysis of *in vivo* perturbations using CED. **A)** LDA UMAP plots showing distribution of single cells from *in vivo* CED experiments expressing the indicated sgRNAs in no radiotherapy or **(B)** radiotherapy conditions. Cells which express sgRNAs other than the one highlighted in color are indicated in gray. **C)** Difference in UMAP gaussian kernel densities between perturbations in radiotherapy (RT) and no treatment (no RT) conditions. Gray bar = mean.

**Figure S6.**
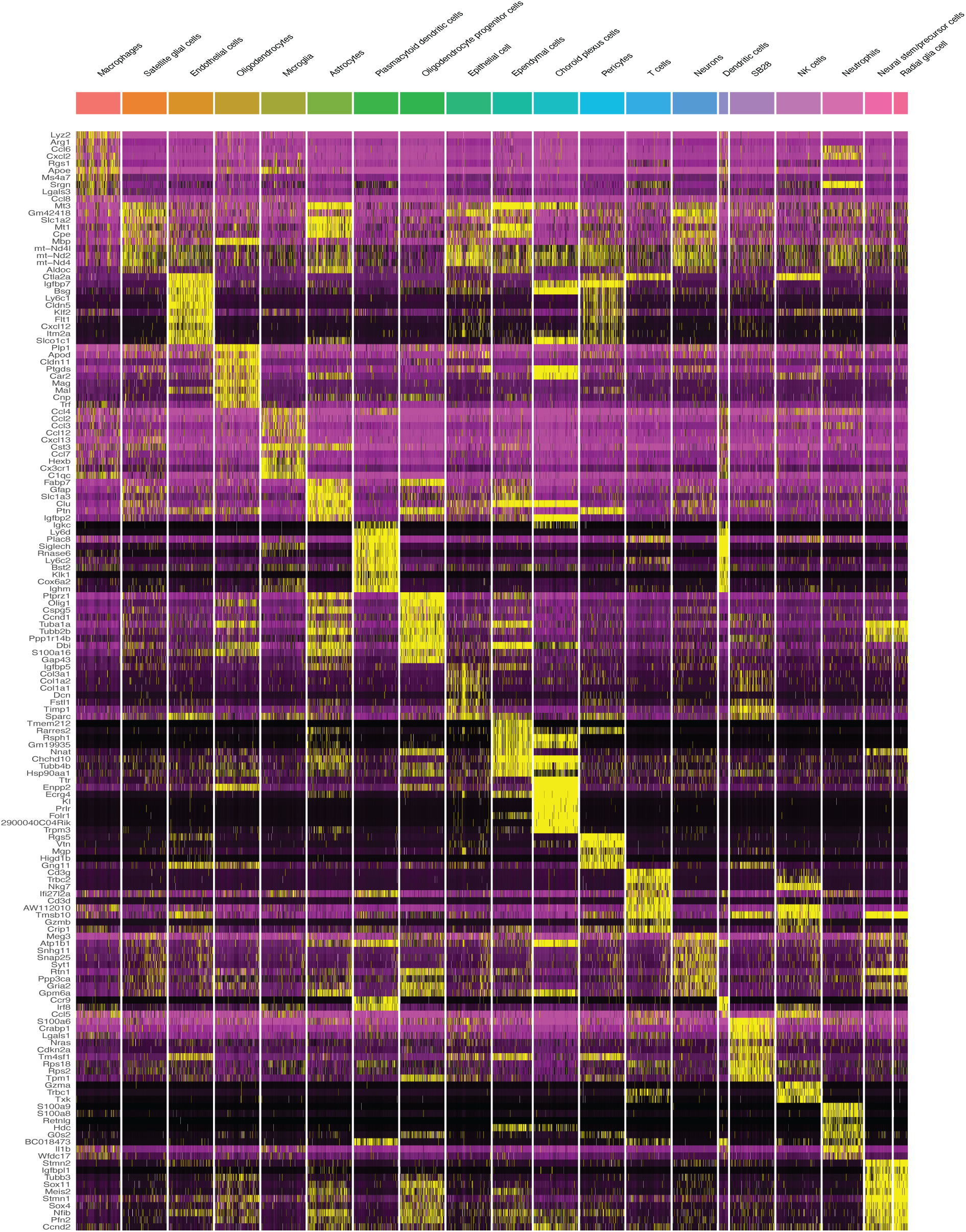
Cell type marker gene expression in SB28 tumors. Heatmap of marker gene expression for each cell type identified by scMRMA with manual annotation of SB28 *in vivo* malignant cells. Expression values represent SCTransform residuals: yellow (maximum) – purple (minimum) colors scale.

**Figure S7.**
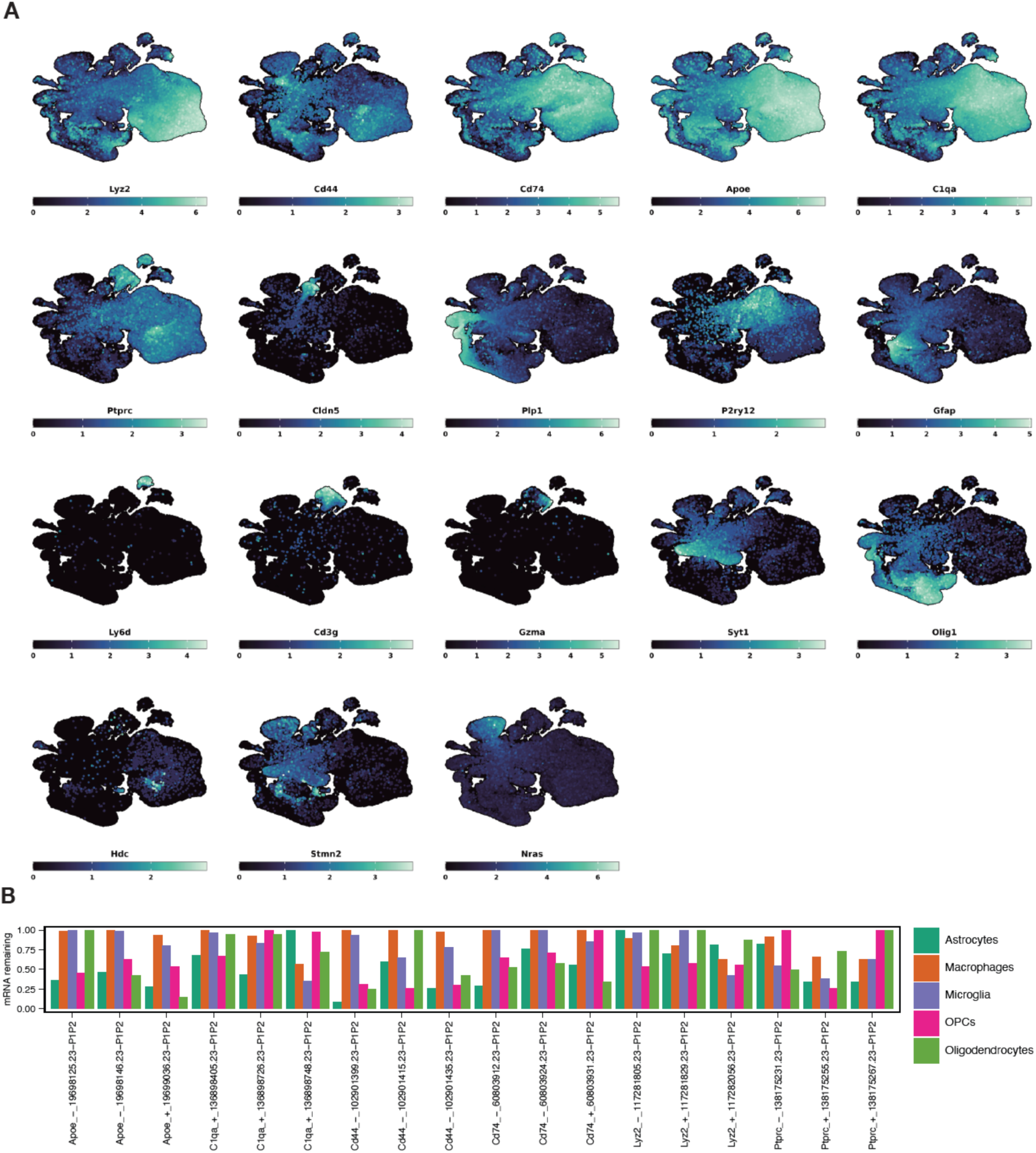
Characterization of SB28 tumors for microenvironment perturbations. **A)** Gene expression featureplots of marker genes or genes selected for sgRNA targeting in SB28 orthotopic tumor cells, overlayed on UMAP coordinates derived in Fig. 5B. Expression values represent log2 corrected UMI’s. **B)** On-target knockdown levels for perturbations of target genes across individual sgRNAs in each major cell type.

